# A Critical Role of Hepatic GABA in The Metabolic Dysfunction and Hyperphagia of Obesity

**DOI:** 10.1101/2020.04.02.022699

**Authors:** Caroline. E. Geisler, Susma. Ghimire, Stephanie. M. Bruggink, Kendra E. Miller, Savanna. N. Weninger, Jason. M. Kronenfeld, Jun. Yoshino, Samuel. Klein, Frank. A. Duca, Benjamin. J. Renquist

## Abstract

Hepatic lipid accumulation is a hallmark of type II diabetes (T2D) and associated with hyperinsulinemia, insulin resistance, and hyperphagia. Hepatic synthesis of GABA, catalyzed by GABA-transaminase (GABA-T), is upregulated in obese mice. To assess the role of hepatic GABA production in obesity-induced metabolic and energy dysregulation, we treated mice with two pharmacologic GABA-T inhibitors and knocked down hepatic GABA-T expression using an antisense oligonucleotide. Hepatic GABA-T inhibition and knockdown decreased basal hyperinsulinemia and hyperglycemia, and improved glucose intolerance. GABA-T knockdown improved insulin sensitivity assessed by hyperinsulinemic-euglycemic clamps in obese mice. Hepatic GABA-T knockdown also decreased food intake and induced weight loss without altering energy expenditure in obese mice. Data from people with obesity support the notion that hepatic GABA production and transport are associated with serum insulin, HOMA-IR, T2D, and BMI. These results support a key role for hepatocyte GABA production in the dysfunctional glucoregulation and feeding behavior associated with obesity.

## Introduction

Type II diabetes (T2D) affects 30 million Americans, while an additional 81 million Americans have pre-diabetes ^1^. Thus, 46% of the U.S. adult population is affected by diabetes, the 7^th^ leading cause of death in America, which consumes 1 in every 7 U.S. health care dollars ^2^. The high prevalence, mortality, and economic burden of T2D underscores a critical need for the development of additional therapeutics to treat diabetes.

Hepatic steatosis is a hallmark of T2D, as accumulation of hepatic lipids strongly correlates with the severity of peripheral insulin resistance and hyperinsulinemia ^3–5^. Furthermore, 87% of diabetics are overweight or obese ^1^. Obesity and T2D are also characterized by dysregulated energy homeostasis, particularly diminished meal-induced satiety which can result in excessive energy intake ^6–8^. Interestingly, hepatic lipid accumulation is directly associated with increased energy intake. In individuals with non-alcoholic fatty liver disease (NAFLD), the percent hepatic steatosis positively correlates with total energy intake and carbohydrate intake ^9^. These findings suggest the possibility that hepatic lipid accumulation may drive obesity induced hyperinsulinemia, insulin resistance, and hyperphagia.

The liver produces and releases hepatokines into circulation in response to acute and chronic nutrient status ^10^. For example, FGF21 and ANGPTL4 are secreted from hepatocytes in response to liver nutrient flux and can act in an endocrine fashion to impact whole body metabolism. In line with this, we have recently established that obesity-induced hepatic lipid accumulation increases hepatocyte production and release of the inhibitory neurotransmitter, GABA, in mice (Geisler et. al co-submitted Fig. 4A) that we hypothesize acts in a paracrine fashion to decrease the firing activity of the hepatic vagal afferent nerve (HVAN) to regulate insulin secretion and sensitivity (Geisler et. al co-submitted Fig. 5). These findings provide a novel mechanistic link explaining how hepatic lipid accumulation, though increasing extracellular hepatocyte-produced GABA to impair HVAN signaling, drives the development of metabolic diseases.

The GABA shunt classically refers to a TCA cycle detour that converts α-ketoglutarate to succinate and concomitantly breaks down a molecule of GABA via GABA-Transaminase (GABA-T) ^11^. However, in the liver, GABA-T mediates GABA synthesis ^12^. We have proposed that hepatic lipids activate reversed GABA shunt activity in hepatocytes, and that hepatic GABA and glucose production are metabolically linked (Geisler et. al co-submitted Fig. 5). It remains completely untested whether manipulating this GABA shunt can prevent hepatic steatosis derived metabolic disease. Thus, GABA-T represents a promising target to decrease hyperinsulinemia and insulin resistance by limiting hepatic GABA production. Accordingly, in the current manuscript, we employed two novel models to limit hepatic GABA production: 1) pharmacologic inhibition of GABA-T activity, and 2) antisense oligonucleotide (ASO) mediated knockdown of hepatic specific GABA-T expression. Using these models for the first time we assessed systemic glucose homeostasis to strengthen the causative role between hepatic GABA production and hyperinsulinemia / insulin resistance. We also assessed food intake and energy expenditure to understand the role of hepatic GABA production in the dysregulation of energy homeostasis in obesity.

## Results

### GABA-Transaminase Inhibition Improves Glucose Homeostasis in Obesity

To directly assess the effect of GABA-T in obesity-induced metabolic dysfunction we treated high fat diet-induced obese mice with one of two irreversible GABA-T inhibitors, ethanolamine-O-sulphate (EOS) or vigabatrin (8 mg/day). Both reduce hepatic GABA-T activity by over 90% within two days ^13^. Through 5 days of treatment, body weight remained similar among EOS, vigabatrin, and saline injected mice (Fig. 1A). Four days of EOS or vigabatrin treatment decreased serum insulin and glucose concentrations and increased the glucose:insulin ratio relative to pre-treatment (Figs. 1B-1D). Two-weeks washout from EOS or vigabatrin resulted in a return of serum insulin and the glucose:insulin ratio to pre-treatment levels (Figs. 1B-1D). EOS treatment (5 days) decreased serum glucagon relative to control mice (Fig. 1E). Glucose clearance during an oral glucose tolerance test (OGTT) was improved by 4 days of GABA-T inhibition (Figs. 1F-1G). Coincident with this improved clearance, glucose stimulated serum insulin was decreased by vigabatrin and tended to be decreased by EOS relative to pre-treatment concentrations (Fig. 1H). Two-weeks washout from EOS and vigabatrin markedly increased glucose stimulated serum insulin (Fig. 1H). Both GABA-T inhibitors improved insulin sensitivity assessed by insulin tolerance test (ITT) within 4 days of initiating treatment (Fig. 1I-1J). As oral drug delivery is preferred in clinic, we conducted another series of studies with EOS provided in the drinking water (3 g/L) for 4 days. EOS in the drinking water similarly improved measures of glucose homeostasis and the response washed out after 2 weeks without EOS in the drinking water (Extended Data Fig. 1).

**Figure 1.**
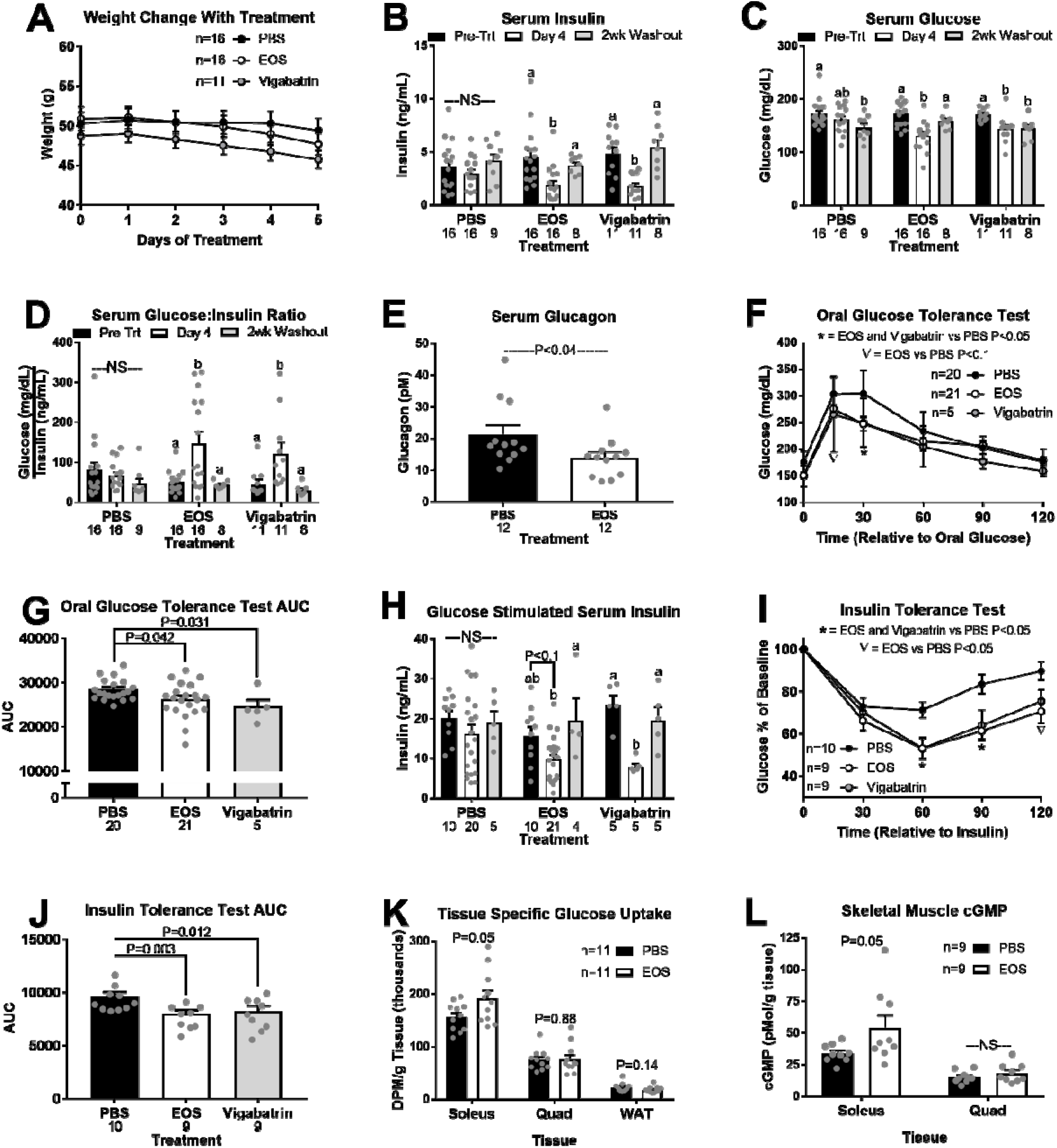
GABA-Transaminase inhibition improves glucose homeostasis in obesity. HFD-induced obes mice were treated with GABA-Transaminase inhibitors ethanolamine-O-sulfate (EOS) or vigabatrin (8mg/day), or phosphate buffered saline (PBS; control) for 5 days. Body weight during treatment (A). Basal serum insulin (B), glucose (C), and glucose:insulin ratio (D) pre-treatment, on treatment day 4, and after a 2-week washout. Serum glucagon in response to EOS (E). Oral glucose tolerance (OGTT; F), and OGTT area under the curve (AUC; G) on treatment day 4. Glucose stimulated serum insulin (H) pre-treatment, on treatment day 4, and after a 2-week washout. Insulin tolerance (ITT; I) and ITT AUC (J) on treatment day 4. Tissue specific ^3^H-2-deoxy-D-glucose (10 μCi/mouse; K) uptake and cGMP content (L) on treatment day 5. DPM = disintegrations per minute, NS = non-significant. ^a,b^ Bars that do not share a common letter differ significantly within injection treatment (P < 0.05; number below bar denotes n per group). All data are presented as mean ± SEM.

To further assess the mechanism by which GABA-T inhibition improves glucose clearance, we measured tissue specific ^3^H-2-deoxy-D-glucose (2DG) uptake following an oral glucose gavage on day 5 of EOS or saline treatment. EOS treatment increased 2DG uptake by the soleus (22%) but did not affect 2DG clearance by the quadriceps femoris (quad) or gonadal white adipose tissue (WAT; Fig. 1K). Given that blood perfusion is a key regulator of insulin action and glucose clearance, we subsequently measured cGMP, a key second messenger downstream of nitric oxide (NO) signaling that regulates blood flow ^14^. EOS increased cGMP in the soleus (59%) but had no effect in quad (Fig. 1L).

In lean mice, which have low hepatic GABA production (Geisler et. al co-submitted Figs. 4A-4B), there was no effect of EOS or vigabatrin (8 mg/day) on serum insulin concentrations, glucose tolerance, or insulin sensitivity (Extended Data Fig. 2). In lean mice, GABA-T inhibition decreased glucose stimulated serum insulin, while EOS decreased serum glucose (Extended Data Figs. 2B and 2F). We propose GABA-T inhibition decreases serum glucose by directly impairing hepatic gluconeogenic flux from TCA cycle intermediates (Geisler et. al co-submitted Fig. 5).

### Hepatic GABA-Transaminase Knockdown Improves Obesity-Induced Metabolic Dysfunction

To overcome the limitation of global pharmacologic inhibitors we next used an ASO model to specifically knockdown hepatic GABA-T expression. Peripherally administered ASOs do not cross the blood brain barrier ^15^. Outside the central nervous system, the liver and pancreas express the most GABA-T ^16^. A GABA-T targeted ASO (12.5 mg/kg IP twice weekly) decreased hepatic GABA-T mRNA expression by > 98% within 1 week. Importantly, this GABA-T targeted ASO did not affect pancreatic or whole-brain GABA-T mRNA expression (Fig. 2A). To establish the key role of GABA-T in liver slice GABA production, we measured *ex vivo* liver slice GABA release. GABA-T knockdown cut obesity induced liver slice GABA release by 61% (Fig. 2B). One week of GABA-T knockdown in obese mice did not affect body weight but decreased basal serum insulin and glucose concentrations and elevated the glucose:insulin ratio (Figs. 2C-2F). GABA-T targeted ASO injections also improved oral glucose clearance without affecting oral glucose stimulated serum insulin concentrations (Figs. 2G-2I), and improved insulin sensitivity compared to scramble control ASO injected mice (Figs. 2J-2K). These findings directly implicate hepatic GABA production in the development of obesity-induced hyperinsulinemia and insulin resistance.

**Figure 2.**
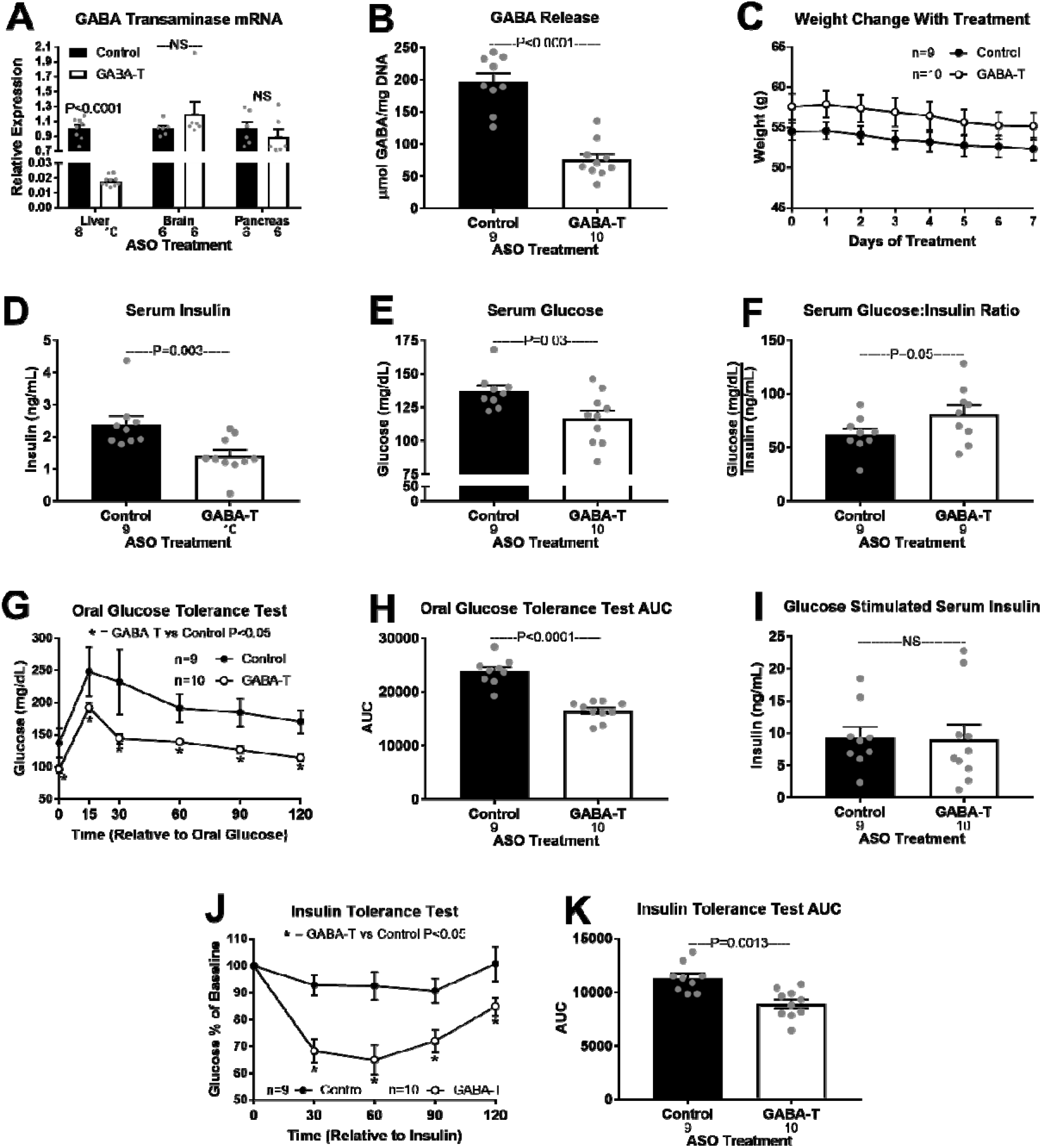
Acute hepatic GABA-Transaminase knockdown improves obesity induced metabolic dysfunction. GABA-T mRNA expression in liver, whole brain, and pancreas after 1 week of injection with a GABA-T targeted or scramble control antisense oligonucleotide (ASO; 12.5 mg/kg IP twice weekly) in high fat diet-induced obese mice (A). Release of GABA (μmol/mg DNA) from hepatic slice (B). Body weight during treatment (C). Basal serum insulin (D), glucose (E), and glucose:insulin ratio (F). Oral glucose tolerance (OGTT; G), OGTT area under the curve (AUC; H), oral glucose stimulated serum insulin (I), insulin tolerance (ITT; J), and ITT AUC (K). Number below bar denotes n per group. NS = non-significant. All data are presented as mean ± SEM.

Compared with control ASO, chronic (4 week) GABA-T ASO treatment decreased *ex vivo* liver slice GABA release, and decreased body mass and serum insulin, while increasing the glucose:insulin ratio, and (Extended Data Fig. 3A-3C, 3E). Thus, the metabolic response to GABA-T knockdown persists. Admittedly, a decrease in body mass with chronic ASO treatment could contribute to the improvements in glucose homeostasis.

To more precisely assess the effect of GABA-T knockdown on insulin action before body mass is affected, we performed hyperinsulinemic-euglycemic clamps in diet-induced obese mice treated with scramble control or GABA-T targeted ASOs for 1 week. Body weight on the day of clamp was not affected by treatment (Fig. 3A). All timepoints are relative to the onset of insulin (4 mU/kg/min) and variable rate 3-[^3^H]-D-glucose in dextrose infusion at time 0, with the basal period referencing pre-0 measurements. Blood glucose concentrations were lower in GABA-T ASO treated mice during the first 20 minutes of the clamp, but not different between ASO treatments from 30-120 minutes of the clamp during which euglycemia was achieved in both groups (Fig. 3B). To maintain the same level of euglycemia, GABA-T knockdown mice required a higher glucose infusion rate (GIR: mg/kg/min) which was significant from 30-120 minutes of the clamp (Fig. 3C). Plasma insulin concentrations did not differ by ASO treatment during either the basal period or during the clamp (Fig. 3D). Importantly, insulin concentrations were elevated during the clamp (Fig. 3D; P=0.013). GABA-T knockdown did not affect the rate of endogenous glucose appearance (Ra) during the basal or clamp periods (Fig. 3E). The basal rate of glucose disappearance (Rd) did not differ between ASO treatments. However, hyperinsulinemia during the clamp nearly doubled Rd in GABA-T knockdown but not control mice (Fig. 3F). After 120 minutes of the clamp, mice received a bolus of ^14^C-2-dexyglucose and were sacrificed 30 minutes later to assess tissue specific glucose uptake. GABA-T knockdown improved glucose uptake by the soleus, quadricep, and calf skeletal muscles (Fig. 3G).

**Figure 3.**
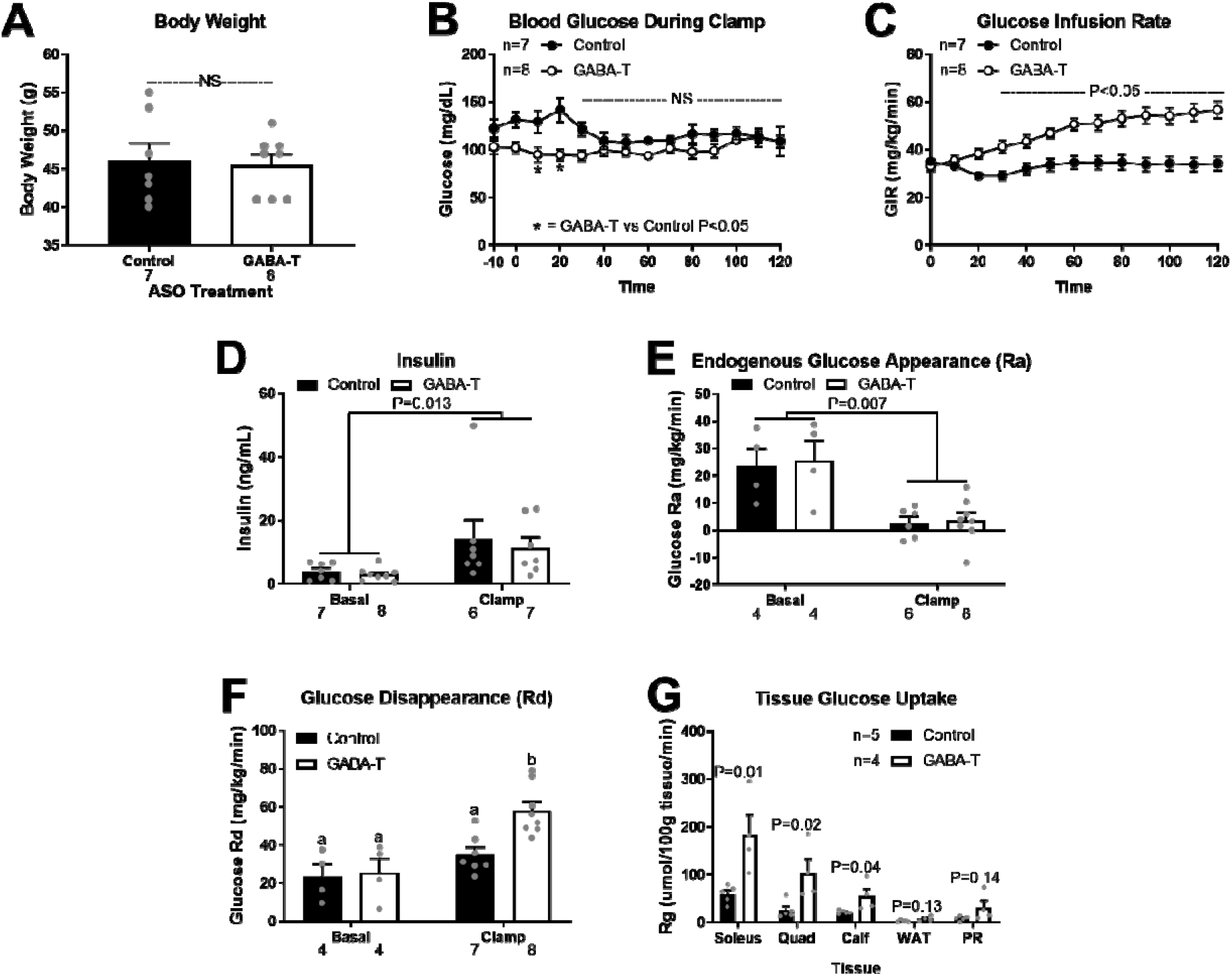
One week of hepatic GABA-Transaminase knockdown improves insulin sensitivity assessed b hyperinsulinemic euglycemic clamp. High fat diet-induced obese mice received 1 week of injections with a GABA-T targeted or scramble control antisense oligonucleotide (ASO; 12.5mg/kg IP twice weekly) before hyperinsulinemic euglycemic clamps were performed. Body weight the day of clamp procedure (A). Blood glucose concentrations (B) and glucose infusion rate during the clamps (C). Serum insulin concentrations before insulin infusion (basal) and during the clamp (D). Endogenous glucose appearance (Ra; E) and glucose disappearance (Rd; F) before insulin infusion (basal) and during the clamp. Tissue specific ^14^C-2-deoxyglucose uptake (G; WAT = white adipose tissue, PR = perirenal adipose tissue). Number below bar denotes n per group. NS = non-significant. All data are presented as mean ± SEM.

In chow fed mice hepatic GABA production is limited (Geisler et. al co-submitted Figs. 4A-4B). In turn, we expected minimal responses to inhibition of the GABA shunt in lean mice. We tested two GABA-T targeted ASOs that both decreased hepatic GABA-T mRNA expression by ≥ 80% within 1 week and ≥ 94% after 4 weeks of treatment. Neither altered pancreatic or whole-brain GABA-T mRNA expression (Extended Data Figs. 4A-4B). Chronic (4 week) GABA-T knockdown did not affect body weight or glucoregulatory measures in lean mice (Extended Data Fig. 4C-4K). This agrees with our observations that acute GABA-T inhibition with EOS does not alter glucose homeostasis in lean mice (Extended Data Fig. 2). Instead, beneficial effects of GABA-T inhibition are specific to obesity, due to an increase in the GABA shunt activity and subsequent hepatic GABA production observed during obesity.

**Figure 4.**
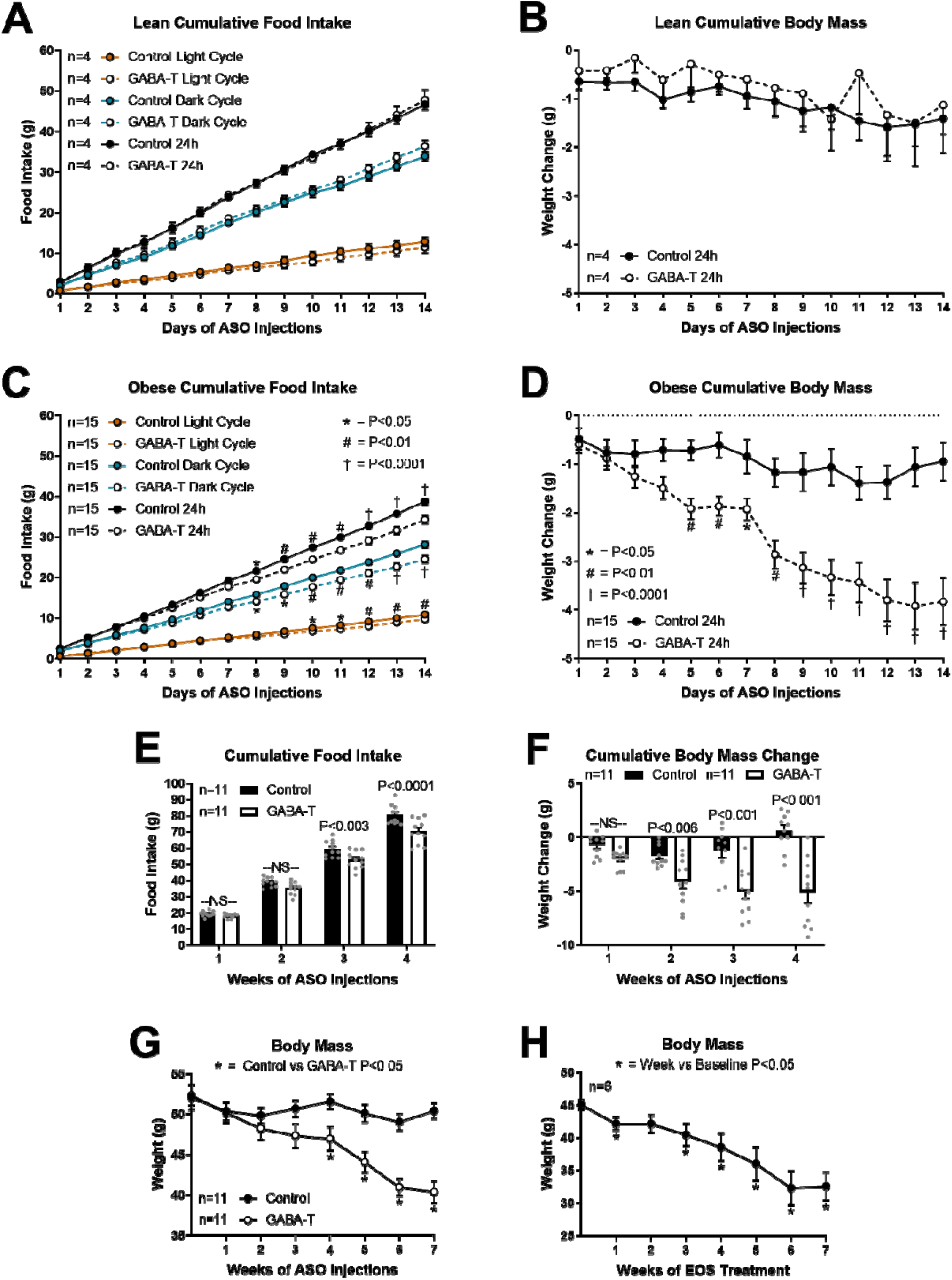
Obesity induced hepatic GABA production increases phagic drive. Cumulative food intake and body mass during the first 2 weeks of GABA-T targeted or scramble control antisense oligonucleotide injections (ASO; 12.5mg/kg IP twice weekly) in lean (A-B) and diet-induced obese (C-G) mice. Cumulative light cycle, dark cycle, and daily food intake (A) and cumulative body mass change (B) in lean mice. Cumulative light cycle, dark cycle, and daily food intake (C) and cumulative body mass change (D) in obese mice. Weekly cumulative food intake (E), cumulative body weight change (F), and body mass (G) during ASO treatment. Body mass during chronic EOS treatment (3 g/L in drinking water). All data are presented as mean ± SEM.

### The Hepatic Vagal Nerve is Essential to Improvements in Glucose Homeostasis Resulting from GABA-Transaminase Inhibition or Knockdown

Hepatic vagotomy prevents the liver from altering afferent signaling to the brain, without affecting normal vagal afferent input originating from the nodose ^17^. In turn, vagotomy prevents inhibition of the vagal afferent tone by GABA produced in the liver. We hypothesize that hepatic GABA mediates the effects that we have reported by acting on the HVAN to induce hyperinsulinemia and insulin resistance. Accordingly, we assessed the effect of EOS treatment in HFD-induced obese hepatic vagotomized and sham operated mice. Although body weight did not differ between surgical groups during EOS treatment (Extended Data Fig. 5A), the response to EOS was dependent on an intact hepatic vagal nerve. In sham operated mice EOS decreased serum insulin and glucose, elevated the glucose:insulin ratio, improved oral glucose tolerance, diminished glucose stimulated serum insulin concentrations, and tended to improve insulin sensitivity as measured by an insulin tolerance test (Extended Data Figs. 5B-5K). Washout restored all parameters to pre-treatment measures. Vagotomy eliminated most of the responses to EOS (Extended Data Figs. 5B-5K). Since hepatic GABA production supports gluconeogenic flux (Geisler et. al co-submitted Fig. 5), we expected GABA-T inhibition to decrease gluconeogenesis through direct actions at the liver. In turn, diminished hepatic glucose output explains the vagal nerve independent decrease in serum glucose during EOS treatment in hepatic vagotomized mice (Extended Data Fig. 5C).

**Figure 5.**
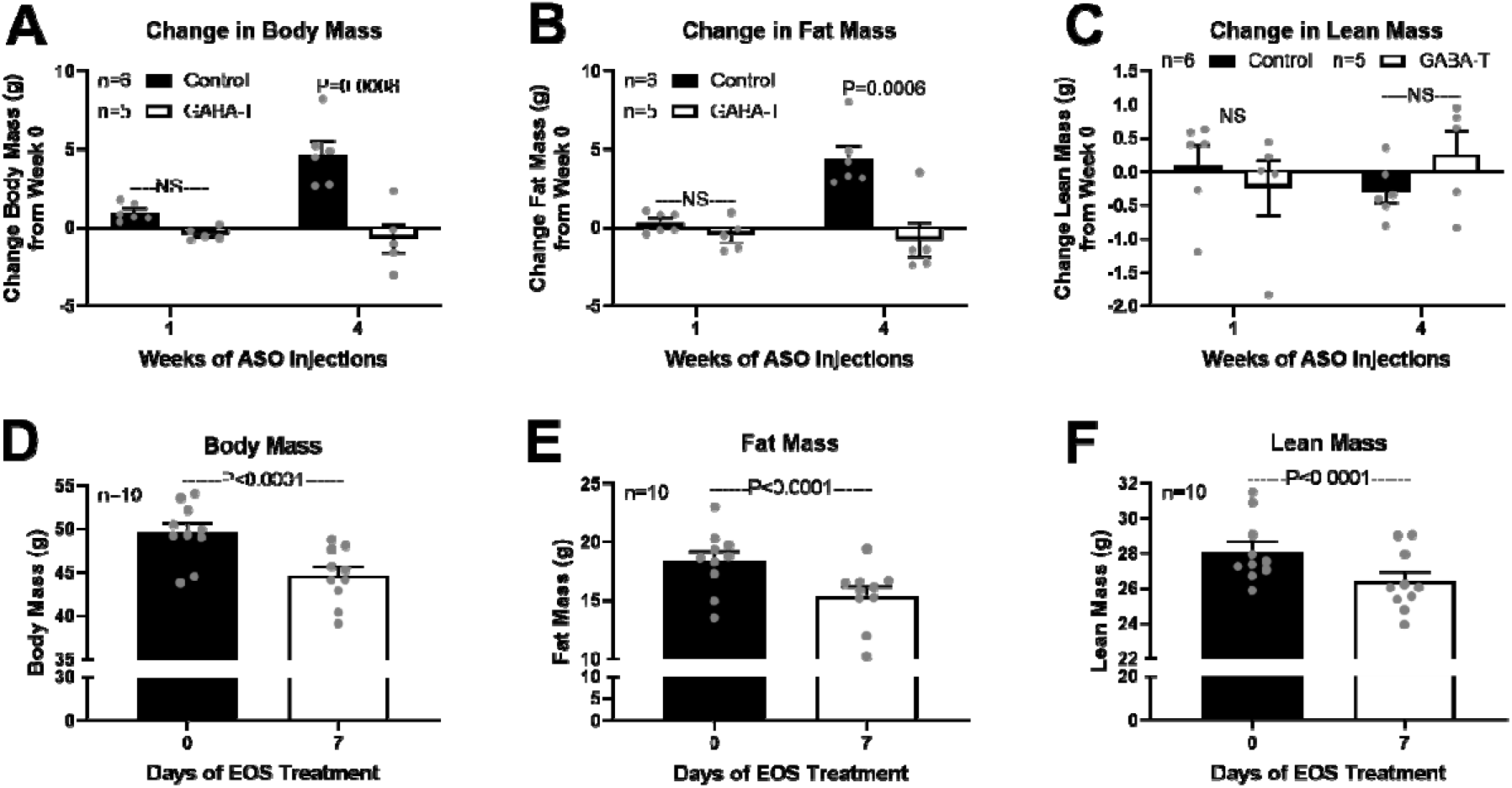
GABA-Transaminase knockdown or inhibition decreases body mass and fat mass. Body composition in antisense oligonucleotide (ASO) treated mice was assessed by Dual-Energy X-ray Absorptiometry (DEXA) at the UCDavis Mouse Metabolic Phenotyping Center. Body composition in ethanolamine-O-sulfate (EOS) treated mice was assessed by EchoMRI 900 at The University of Arizona. Change in body mass (A), fat mass (B), and lean mass (C) after 1 and 4 weeks of GABA-T targeted or scramble control ASO (12.5 mg/kg IP twice weekly) relative to pre-treatment body composition. Body mass (D), fat mass (E), and lean mass (F) on day 0 and 7 of EOS treatment (3 g/L in drinking water). All data are presented as mean ± SEM.

We next assessed the effect of 4 weeks of GABA-T knockdown (12.5 mg/kg GABA-T targeted ASO IP twice weekly) in diet-induced obese sham operated and hepatic vagotomized mice. Consistent with previous observations (Geisler et. al co-submitted Fig. 1B), pre-treatment body weight was lower in obese, vagotomized mice compared to sham operated control mice (Extended Data Fig. 6A). As shown with pharmacological GABA-T inhibition (EOS), GABA-T knockdown decreased basal serum glucose concentrations independent of the hepatic vagal nerve (Extended Data Fig. 6C), again likely due to reduced hepatic glucose output. GABA-T knockdown decreased basal serum insulin, improved oral glucose tolerance, limited oral glucose stimulated serum insulin, and improved insulin sensitivity in sham, but not in vagotomized mice (Extended Data Figs. 6B, 6E-J). Hepatic vagotomy protects against obesity-induced insulin resistance (Geisler et. al co-submitted Fig. 1). Here we show that GABA-T knockdown allowed sham operated animals to achieve similar glucose tolerance, glucose stimulated serum insulin, and insulin sensitivity to that measured in hepatic vagotomized mice.

### GABA-T Knockdown Decreases Fat Mass by Decreasing Food Intake Without Affecting Energy Expenditure in Obesity

Given that hepatic GABA-T knockdown for 4 weeks decreased body weight in obese mice (Extended Data Fig. 3B), we hypothesized that GABA-T knockdown may decrease food intake or increase energy expenditure. Accordingly, we assessed daily food intake and body mass during the light and dark cycle for the first 2 weeks of ASO treatment in lean and obese mice. In lean mice, GABA-T knockdown did not affect cumulative light cycle, dark cycle, or daily food intake (Fig. 4A). Cumulative daily body mass change was also not affected by GABA-T knockdown in lean mice (Fig. 4B). In obese mice, GABA-T knockdown decreased cumulative light cycle, dark cycle, and daily food intake (Fig. 4C).

Cumulative weekly food intake remained lower through 4 weeks of GABA-T knockdown (Fig. 4E). This suppression of food intake was accompanied by a decrease in body mass in GABA-T ASO treated mice (Fig. 4D and 4F). In fact, GABA-T knockdown continued to decrease body weight during 7 weeks of treatment (Fig. 4G). Similarly, 7 weeks of continuous exposure to EOS in the drinking water (3 g/L) dramatically decreased body mass in diet-induced obese mice (Fig. 4H). These data suggest that sensitivity to the weight loss effect of limiting hepatocyte GABA production is maintained throughout treatment but is less significant as body mass returns to normal.

Interestingly, GABA-T knockdown did not alter food intake in response to a 16-hour fast in either lean or obese mice (Extended Data Figs. 7A-7B). This proposes that hepatic GABA signals to regulate *ad libitum* light and dark cycle food intake without affecting the fasting-induced drive for refeeding. Accordingly, GABA-T knockdown did not affect fasting mRNA expression of the canonical hypothalamic neuropeptides regulating food intake (NPY, AgRP, and POMC; Extended Data Fig. 7C). Admittedly, we cannot rule out an effect of central GABA-T knockdown as we did observe a 14% reduction in hypothalamic GABA-T expression at 7 weeks of GABA-T ASO injections (Extended Data Fig. 7C). The anorexigenic hormone leptin induces satiety and weight loss, while leptin resistance in obesity contributes to hyperphagia and weight gain ^18,19^. We tested whether GABA-T knockdown in obesity may have improved leptin sensitivity as a potential mechanism to decrease appetite and cause weight loss. A single 6AM injection of leptin (2 mg/kg) did not affect food intake in mice on either ASO treatment at any timepoint, suggesting that mice on both treatments remained leptin resistant (Extended Data Fig. 7D). Consistent with the decreased food intake previously observed in response to GABA-T knockdown, we again saw that GABA-T knockdown decreased light cycle, dark cycle, and 24-hour food intake independent of injection (Extended Data Fig. 7D). Relative weight change was also not affected by leptin in control or GABA-T ASO obese mice (Extended Data Fig. 7E). Thus, enhanced leptin sensitivity is not mediating the diminished phagic drive in response to GABA-T knockdown in obesity. A single 6AM leptin injection decreased light cycle and 24-hour food intake in chow fed mice without affecting body weight (Extended Data Figs. 7F-7G).

We next investigated whether altered energy expenditure contributed to the body mass loss with GABA-T knockdown in obesity. Energy expenditure (kcal/hr), assessed by respiration calorimetry and corrected using the covariate of lean body mass, was not altered by 1 or 4 weeks of GABA-T knockdown (Extended Data Figs. 8A-8C). In addition, there was no effect of GABA-T knockdown on the composition of oxidized macronutrients (respiratory exchange ratio; RER), daily water intake, or daily activity (Extended Data Figs. 8D-8I). Thus, the effects of GABA-T knockdown on body mass appear independent of energy expenditure or nutrient oxidation.

Body composition, assessed by Dual-Energy X-ray Absorptiometry (DEXA), revealed that 4 weeks of GABA-T knockdown decreased total body mass and fat mass without affecting lean mass (Figs. 5A-5C). The decreased fat mass with GABA-T knockdown suggests that weight loss induced by limiting hepatocyte GABA production reflects a loss of adiposity. We additionally assessed body composition on day 0 and 7 of providing obese mice with EOS in their drinking water (3 g/L). We found that EOS treatment decreased total body mass (10%), fat mass (16.27%), and lean mass (6.15%). Although there was a small loss of lean mass, diminished fat mass contributes the majority of lost body mass with EOS treatment.

### Hepatic Vagotomy Decreases Light Cycle Food Intake on HFD, while GABA-Transaminase Knockdown Normalizes Sham Mice Food Intake to Vagotomy Mice

We had previously reported that hepatic vagotomy decreased weight gain on a high fat diet (Geisler et. al co-submitted Fig. 1B). Here we show that hepatic vagotomy decreases 1-week cumulative light cycle food intake by 22% in diet-induced obese mice, resulting in a 5.3% decrease in cumulative 24-hour food intake (Extended Data Fig. 9A). We continued daily food intake measurements for the next 2 weeks as all mice were treated with the GABA-T targeted ASO. In response to GABA-T knockdown, the previously observed difference in food intake in sham operated and vagotomized mice was eliminated. In fact, cumulative light cycle, dark cycle, and daily food intake were similar in sham operated and vagotomized mice (Extended Data Fig. 9B). The GABA-T ASO resulted in a greater cumulative week 4 body mass loss in sham operated than in vagotomized mice which experienced no net change in body mass (Extended Data Fig. 9D). Thus, the glucoregulatory, phagic, and body mass changes in response to GABA-T knockdown all appear to be dependent on the hepatic vagal afferent nerve. These data support that hepatic vagotomy eliminates the GABA-induced decrease in HVAN activity from being communicated to the brain and accordingly protects against obesity-induced perturbations of glucose and energy homeostasis.

### Associations Between Hepatic GABA System and Glucoregulatory Markers in People with Obesity

To understand the potential clinical relevance of these findings, we assessed the hepatic mRNA expression of *ABAT* (encodes for GABA-T) and GABA transporter (*SLC6A6,* encodes for taurine transporter, TauT; *SLC6A8;* encodes for the creatine transporter, CRT; *SLC6A12;* encodes for the Betaine-GABA Transporter 1, BGT1, and *SLC6A13*; encodes for GABA Transporter 2, GAT2) in 19 people with obesity (age 45 ± 3 yrs old, 2 men and 17 women) who were carefully characterized by measuring intrahepatic triglyceride (IHTG) content using magnetic resonance imaging (MRI) and insulin sensitivity using the hyperinsulinemic-euglycemic clamp procedure (HECP) in conjunction with stable isotopically labeled glucose tracer infusion). The subjects had a wide range in IHTG content, plasma insulin concentration and measures of insulin sensitivity (Extended Data Table 1). There was a trend toward a positive correlation between IHTG content and basal plasma insulin concentration (P=0.06) and a negative correlation between IHTG content and hepatic insulin sensitivity index (HISI, a product of the basal endogenous glucose production rate and plasma insulin concentration, P=0.07; Extended Data Table 2).

Hepatic *ABAT* mRNA expression was positively associated with plasma insulin and negatively with HISI (Figs. 6A-6B). Analysis of single nucleotide polymorphisms (SNPs) reported on the Accelerating Medicines Partnership Type 2 Diabetes Knowledge Portal database shows that SNPs in the reporter region of *ABAT* are associated with decreased risk of T2D (Fig 6C; Extended Data Table 3). Importantly, we found no promoter region SNPs that were significantly associated with an increased odds ratio for T2D. Moreover, given the key role of GABA-T in the central nervous system it is not surprising that SNPs in the *ABAT* gene were not identified. These data support the notion that GABA-T is a driver of insulin resistance in people.

**Figure 6.**
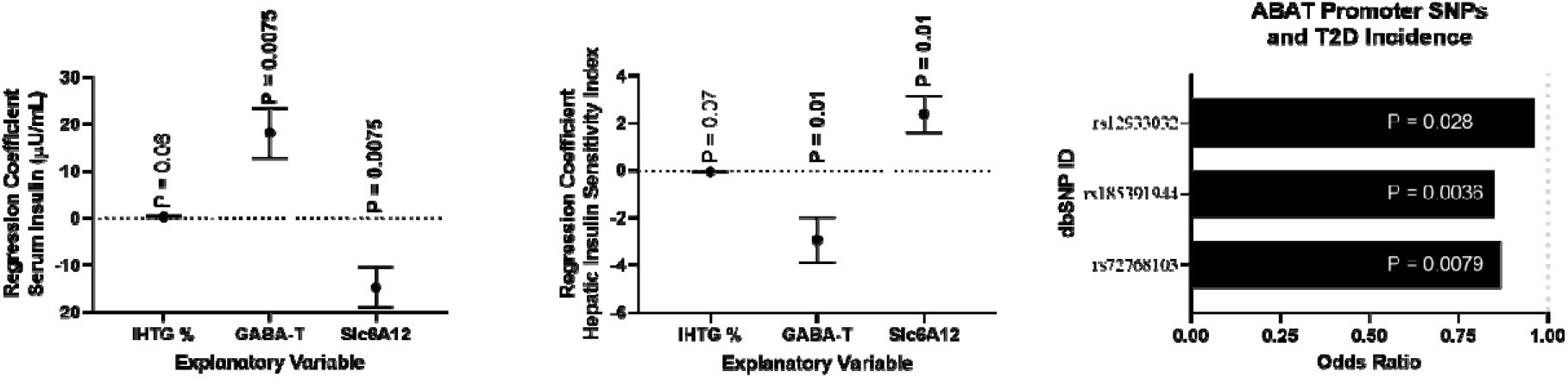
Associations between hepatic GABA system and glucoregulatory markers in people wit obesity. Multivariate regressions including intrahepatic triglyceride % (IHTG%), hepatic *ABAT* (GABA-T) mRNA, and the hepatic GABA transporter (*SLC6A12*) mRNA as explanatory variables for variation in serum insulin (A) or hepatic insulin sensitivity index (HISI; B), *ABAT* and *SLC6A12* mRNA (FPKMUQ; fragments per kilobase million reads upper quartile) were quantified by RNA-Seq from liver tissue. Single nucleotide polymorphisms (SNPs) in the *ABAT* promoter are associated with a decreased risk of type 2 diabetes (T2D; E). All data are presented as mean ± SEM.

By using our *ex vivo* liver slice model we have shown that pharmacologically inhibiting BGT1 increased media GABA content, proposing that although this transporter is bidirectional, flux is more heavily driven towards GABA import (Geisler et al. co-submission Fig. 5). Moreover, missense mutations in *SLC6A12* are associated with an increased risk of T2DM. In fact, a premature stop codon is associated with a 16-fold increase in the risk for T2DM (Geisler et al. co-submission Extended Data Table 3). Here we show that hepatic *SLC6A12* mRNA is negatively associated with plasma insulin concentrations and positively associated with HISI (Figs. 6A and 6B). Thus, we hypothesize that *SLC6A12* expression is primarily indicative of GABA re-uptake into hepatocytes, which depresses GABA signaling onto the HVAN.

Since we have established that GABA-T knockdown and inhibition reduces food intake and body weight, we utilized the knowledge portal diabetes database to identify SNPs associated with BMI. We found missense mutations in *SLC6A12* (6), *SLC6A6* (3), and *SLC6A13* (4) that were associated with increased BMI (Extended Data Table 4). Taken together, the data obtained from the studies conducted in people support the potential clinical translation of our findings in the mouse.

## Discussion

The liver signals in an endocrine fashion through a myriad of hepatokines ^10^. Many hepatokines change with the degree of hepatic steatosis and are linked to altered metabolic homeostasis. We are adding liver produced GABA to the list of steatosis affected hepatokines that alter glucose homeostasis. Unlike other hepatokines, which act in an endocrine fashion, GABA is acting locally in a paracrine fashion on the parasympathetic nervous system to mediate its downstream metabolic effects. Muting hepatic GABA production by pharmacologic inhibition or ASO mediated knockdown of GABA-T attenuates obesity-induced hyperinsulinemia and insulin resistance. Beyond the improvements in glucose homeostasis, hepatic GABA-T knockdown decreases food intake causing a decrease in adiposity and weight loss.

### Glucose Homeostasis

Hepatic GABA production improves insulin sensitivity primarily by increasing skeletal muscle glucose clearance (Figs. 1K and 3G). Our results propose that some of the improvements in glucose clearance may be mediated by increased blood flow. Vasodilation of the microvasculature accounts for 40% of insulin stimulated muscle glucose uptake ^20^. Insulin and acetylcholine signaling at endothelial cells stimulates NO production, which signals to smooth muscle cells inducing cGMP production and vasodilation ^14,21^. The microvascular vasodilatory response is reduced in obesity, directly contributing to systemic insulin resistance ^22–24^. We showed that EOS treatment increased soleus muscle cGMP concentrations (Fig. 1L). Future work will focus on understanding how hepatic GABA signaling at the HVAN regulates endothelial NO production and muscle perfusion.

### Food Intake Regulation

Although weight loss itself improves insulin sensitivity and metabolic health ^25^, GABA-T knockdown improves glucose homeostasis independent of the decrease in food intake and body weight. One week of GABA-T ASO treatment does not decrease food intake (Fig. 4E) or body weight (Figs. 2C and 4F) compared to controls, yet basal serum insulin concentrations, glucose tolerance, and insulin sensitivity are markedly improved (Figs. 2D, 2G, and 2J).

The HVAN has long been implicated in transmitting liver derived signals to the hindbrain to regulate feeding behavior ^26,27^. Early work established that hepatic portal infusions of glucose, amino acids, and lipids suppress food intake ^28–31^, while more recent studies support that carbohydrate signals originating from the liver regulate feeding behavior through HVAN dependent mechanisms ^32,33^. Peripheral satiation factors including glucagon like peptide-1 (GLP-1), cholecystokinin (CCK), lipids, and leptin all reduce food intake dependent upon increasing gastric and hepatic vagal branch afferent firing, while the orexigenic hormone ghrelin suppresses vagal afferent tone ^29,34–38^. We propose a novel addition to this regulation of vagal nerve activity by metabolites and gut hormones, suggesting that hepatic lipid accumulation stimulates hepatic GABA release to depress HVAN activity. Consistent with the effects of other appetite regulators, GABA mediated suppression of HVAN activity would be expected to increase phagic drive, while removal of this inhibitory signal would be expected to increase HVAN firing and reduce food intake. In lean mice, which have low levels of hepatic GABA release, GABA-T knockdown did not alter food intake. Thus, hepatocyte released GABA represents a novel orexigenic signal enhanced by obesity.

Diet-induced obesity dysregulates the diurnal pattern of food intake. Mice on a chow diet eat ~20% of their daily food intake during the light cycle while this increases to ~30% with HFD feeding ^39^. In healthy mice, vagal afferent receptor populations are regulated by nutritional state, expressing an orexigenic profile during fasting and an anorexigenic profile upon refeeding ^40^. In turn, in chow fed mice, anorexigenic mechano-stimuli more effectively stimulate gastric vagal afferent nerve activity during the light cycle ^41^. These circadian patterns are lost with diet-induced obesity ^40,42^. Together these changes promote increased food intake during the light cycle and support the hypothesis that the dysregulation of feeding behavior in obesity is partially mediated by aberrant vagal signaling.

In lean mice, hepatic vagotomy shifts the diurnal feeding pattern to increase light cycle food intake ^43,44^, potentially mediated by the loss of peripheral light cycle anorexigenic stimuli. We report that diet-induced obese, hepatic vagotomized mice eat less during the light cycle than sham operated controls, suggesting that hepatic vagotomy protects against the obesity-induced increase in daytime feeding. We hypothesize that preventing the GABA mediated depression of HVAN activity from reaching the hindbrain not only improves glycemic control (Geisler et. al co-submitted Fig. 1) but decreases light cycle food intake, explaining the decreased weight gain with HFD feeding in hepatic vagotomized mice (Geisler et. al co-submitted Fig. 1B). Further supporting a role of hepatic GABA in HFD induced weight gain, we previously observed that inducing hepatic Kir2.1 expression limited hepatic GABA release and HFD induced weight gain (Geisler et. al co-submitted Fig. 3C).

### Conclusion

Weight loss by people with obesity is often difficult to achieve and maintain. It is likely that dysregulated satiety signaling impairs the effectiveness of weight loss strategies. This work identifies hepatic GABA production as a potential therapeutic target which independently improves systemic glucose homeostasis and decreases food intake in obesity. In people with obesity, hepatic GABA production and transport are associated with glucoregulatory markers, T2D, and BMI supporting the potential translation of this work to improve metabolic health in people.

## Extended Data Titles and Legends

**Extended Data Figure 1.**
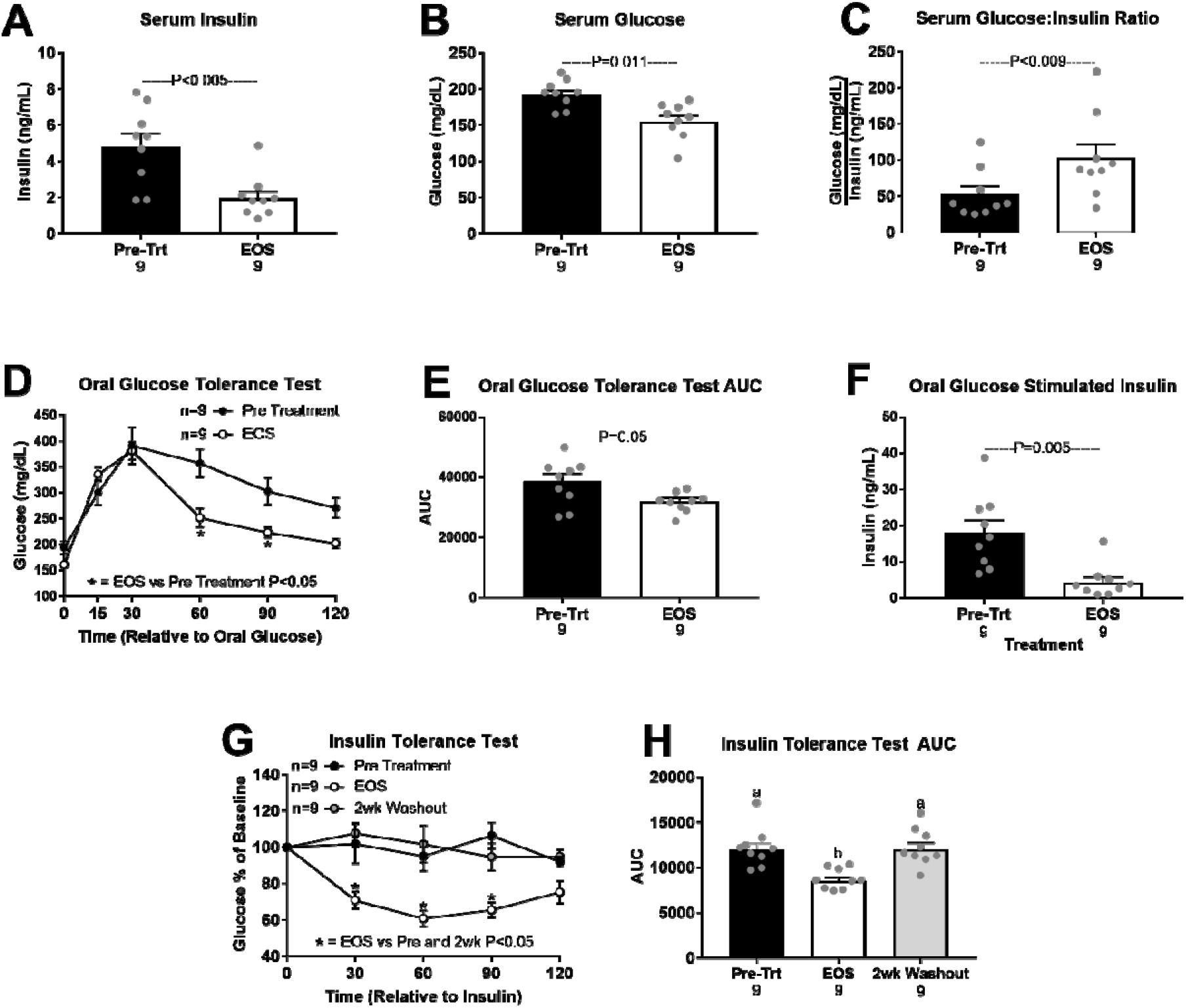
Glucose homeostasis in obese male mice treated with the GABA-Transaminase inhibitor ethanolamine-O-sulfate (EOS; 3 g/L in drinking water). EOS effects on serum insulin (A) glucose (B), and glucose:insulin ratio (C) pre-treatment and after 4 days of treatment. Oral glucose tolerance (OGTT; D) OGTT area under the curve (OGTT AUC; E), and oral glucose stimulated insulin (F) pre-treatment and after 3 days of treatment. Insulin tolerance (ITT; G) and ITT AUC (H) pre-treatment, on day 4 of treatment (EOS), and after a 2-week washout period. ^a,b^ Bars that do not share a common letter differ significantly (P < 0.05; number below bar denotes n per group). All data are presented as mean ± SEM.

**Extended Data Figure 2.**
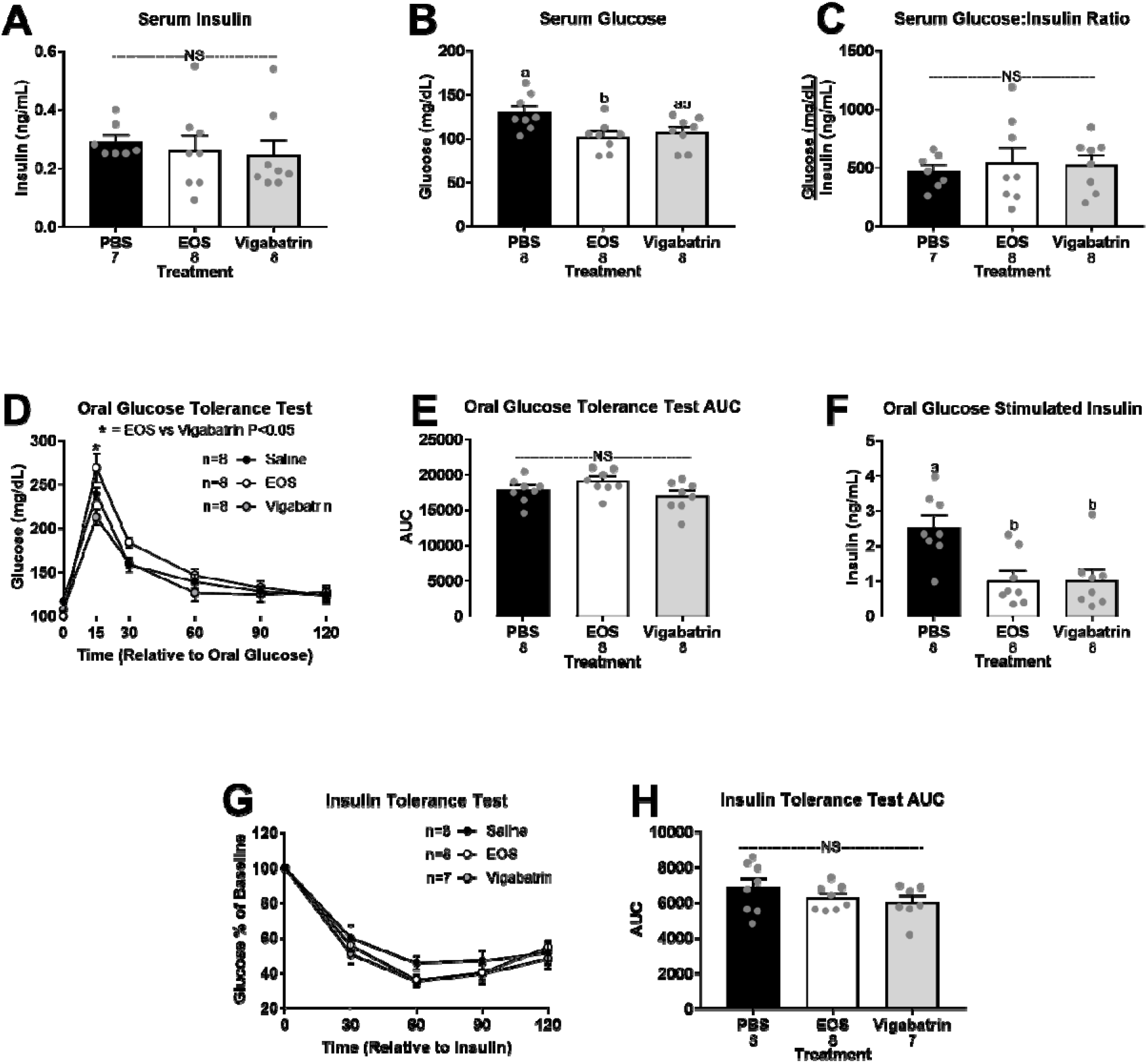
Glucose homeostasis in lean male mice treated with GABA-Transaminase inhibitors ethanolamine-O-sulfate (EOS) or vigabatrin (8 mg/day), or phosphate buffered saline (PBS; control). Serum insulin (A), glucose (B), and glucose:insulin ratio (C) on treatment day 4. Oral glucose tolerance (OGTT; D), OGTT area under the curve (AUC; E), and oral glucose stimulated serum insulin (F) on treatment day 3. Insulin tolerance (ITT; G) and ITT AUC (H) on treatment day 4. NS = non-significant. ^a,b^ Bars that do not share a common letter differ significantly (P < 0.05; number below bar denotes n per group). All data are presented as mean ± SEM.

**Extended Data Figure 3.**
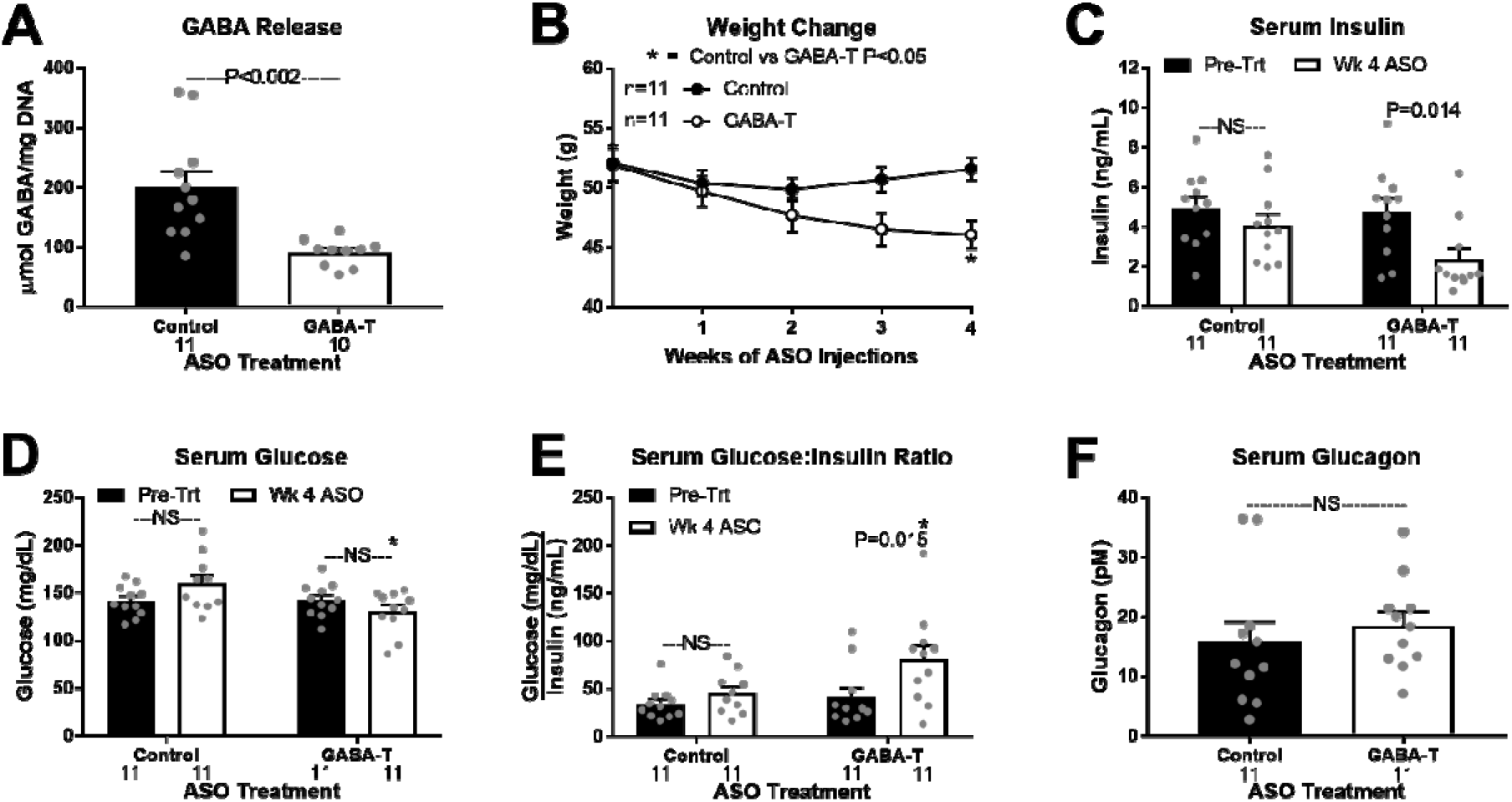
Chronic hepatic GABA-Transaminase knockdown improves obesity induced metabolic dysfunction. High fat diet-induced obese mice were treated for 4 weeks with a GABA-T targeted or scramble control antisense oligonucleotide (ASO; 12.5 mg/kg IP twice weekly). Release of GABA (μmol/mg DNA) from hepatic slices (A). Body weight during treatment (B). Basal serum insulin (C), glucose (D), and glucose:insulin ratio (E) pre-treatment and after 4 weeks of treatment. Serum glucagon (F) after 4 weeks of treatment. Number below bar denotes n per group. NS = non-significant. All data are presented as mean ± SEM.

**Extended Data Figure 4.**
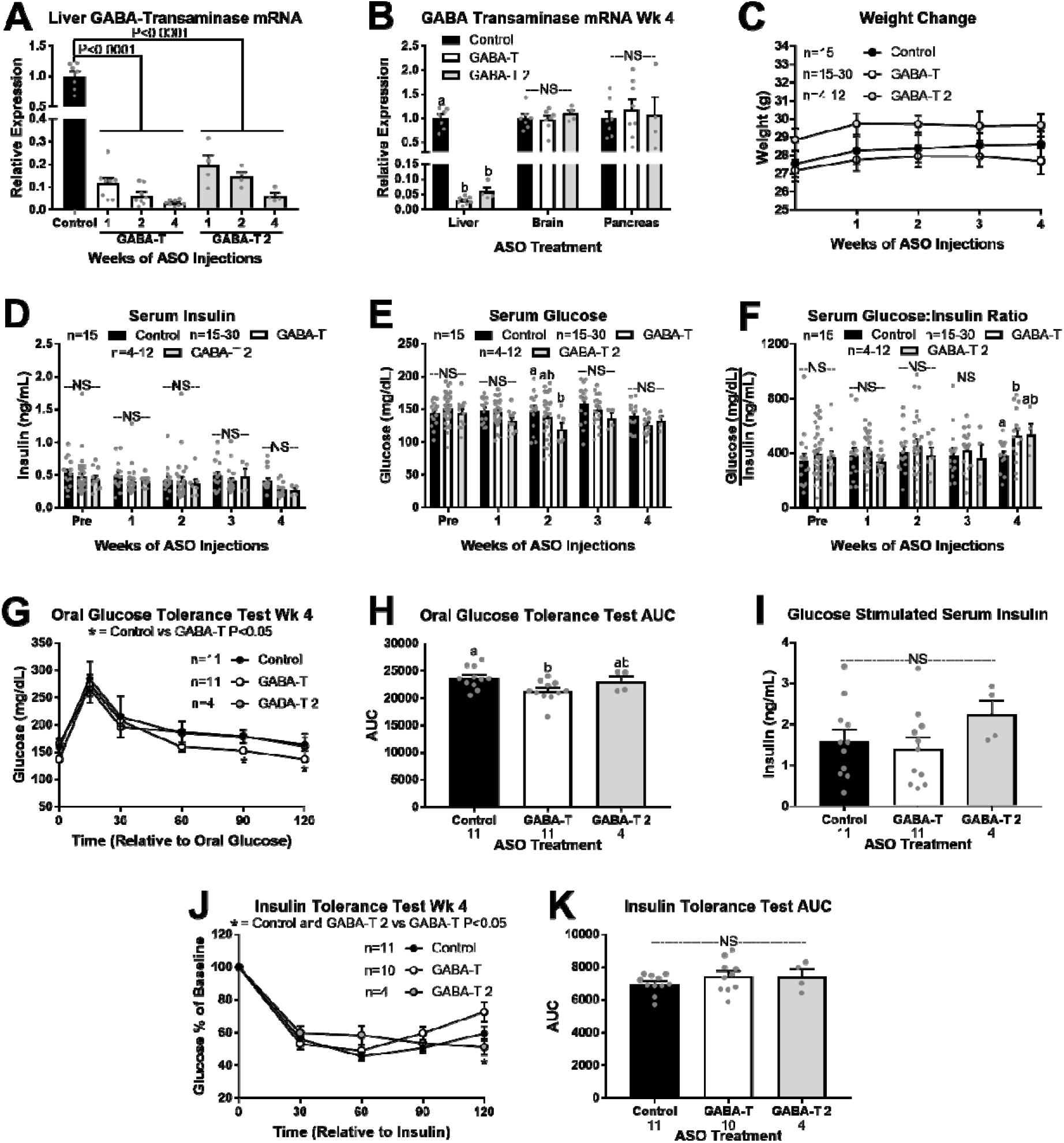
Glucose homeostasis in lean mice treated with the scramble control antisense oligonucleotide (ASO), or 1 of 2 GABA-Transaminase (GABA-T) targeted ASO sequences (GABA-T or GABA-T 2; 12.5 mg/kg IP twice weekly) for 4 weeks. Hepatic GABA-T mRNA expression after 1, 2, and 4 weeks of ASO injections (A). GABA-T mRNA expression in liver, whole-brain, and pancreas after 4 weeks of ASO injections (B). Body weight during treatment (C). Basal serum insulin (D), glucose (E), and glucose:insulin ratio (F) pre-treatment and after 1, 2, 3, and 4 weeks of treatment. Oral glucose tolerance (OGTT; G), OGTT area under the curve (AUC; H), oral glucose stimulated serum insulin (I), insulin tolerance (ITT; J), and ITT AUC (K). ^a,b,c^ Bars that do not share a common letter differ significantly (P < 0.05). Number below bar denotes n per group. NS = non-significant. All data are presented as mean ± SEM.

**Extended Data Figure 5.**
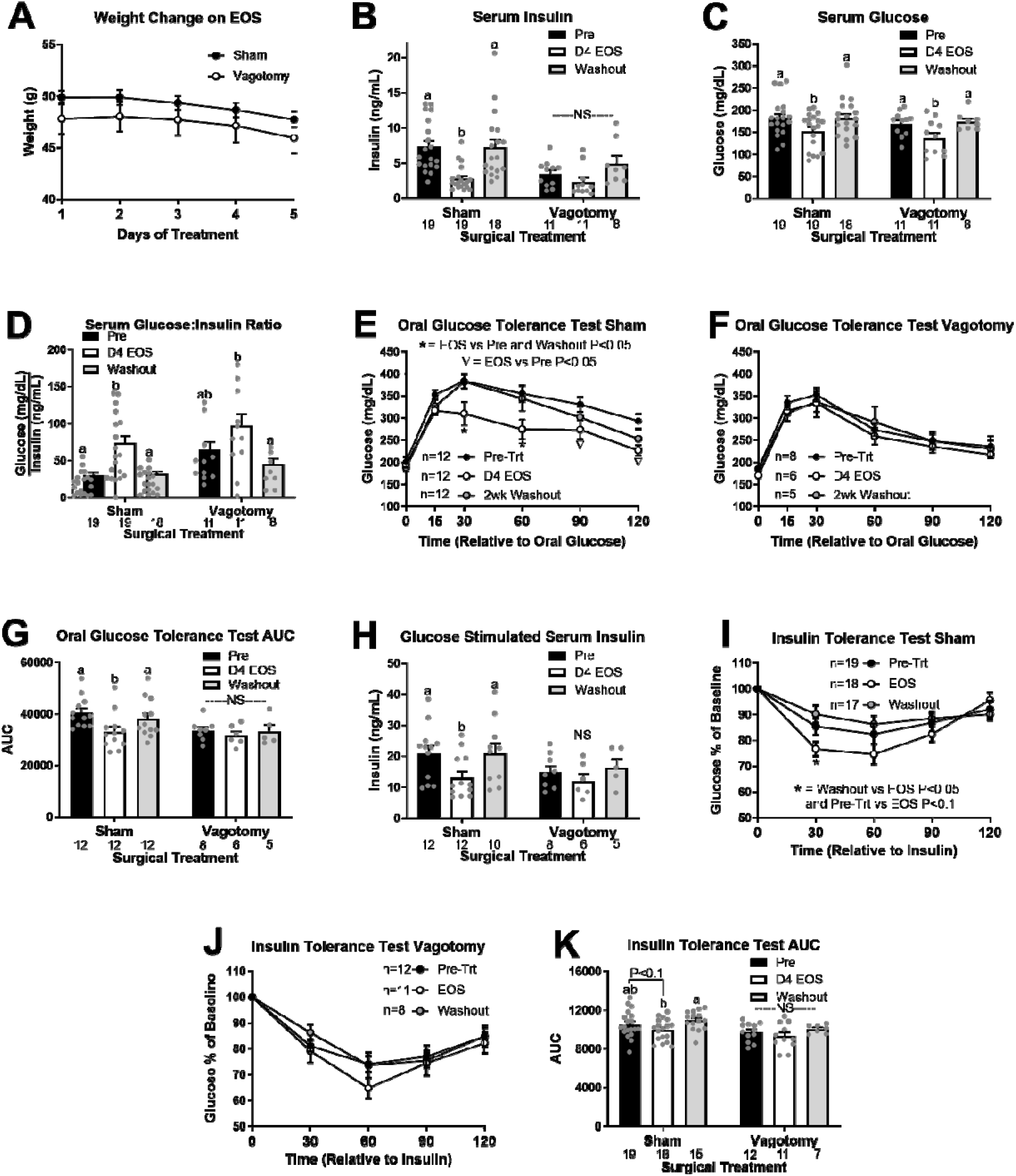
GABA-Transaminase inhibition improves glucose homeostasis in sham but not vagotomy mice. HFD induced sham operated and hepatic vagotomized mice were treated with the GABA-Transaminase inhibitor ethanolamine-O-sulfate (EOS) (8mg/day) for 5 days. Body weight during treatment (A). Basal serum insulin (B), glucose (C), and glucose:insulin ratio (D) pre-treatment, on treatment day 5, and after a 2-week washout. Oral glucose tolerance in sham mice (OGTT; E), oral glucose tolerance in vagotomized mice (F) OGTT area under the curve (AUC; G), and glucose stimulated serum insulin (H) pre-treatment, on treatment day 4, and after a 2-week washout. Insulin tolerance in sham mice (ITT; I) and vagotomized mice (J), and ITT AUC (K) at pre-treatment, on treatment day 5, and after a 2-week washout. NS = non-significant. ^a,b^ Bars that do not share a common letter differ significantly within injection treatment (P < 0.05; number below bar denotes n per group). All data are presented as mean ± SEM.

**Extended Data Figure 6.**
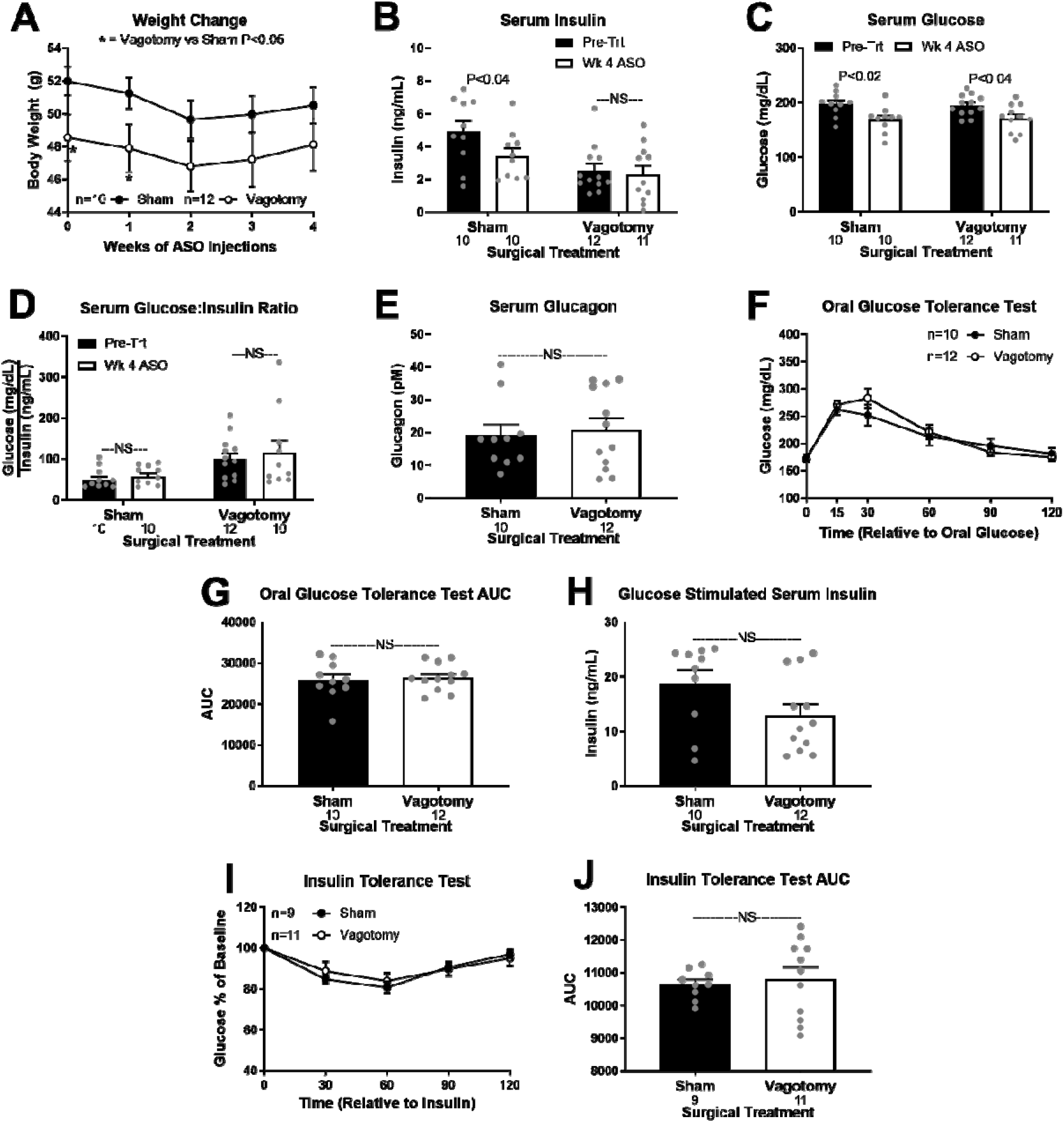
Hepatic GABA-Transaminase knockdown mediated improvements in glucose homeostasis are dependent on an intact hepatic vagal nerve. Diet-induced obese hepatic vagotomized and sham operated mice were treated with a GABA-T targeted antisense oligonucleotide (ASO; 12.5 mg/kg IP twice weekly) for 4 weeks. Body weight during treatment (A). Basal serum insulin (B), glucose (C), and glucose:insulin ratio (D) pre-treatment and after 4 weeks of treatment. Serum glucagon (E), oral glucose tolerance (OGTT; F), OGTT area under the curve (AUC; G), oral glucose stimulated serum insulin (H), insulin tolerance (ITT; I), and ITT AUC (J) after 4 weeks of treatment. Number below bar denotes n per group. NS = non-significant. All data are presented as mean ± SEM.

**Extended Data Figure 7.**
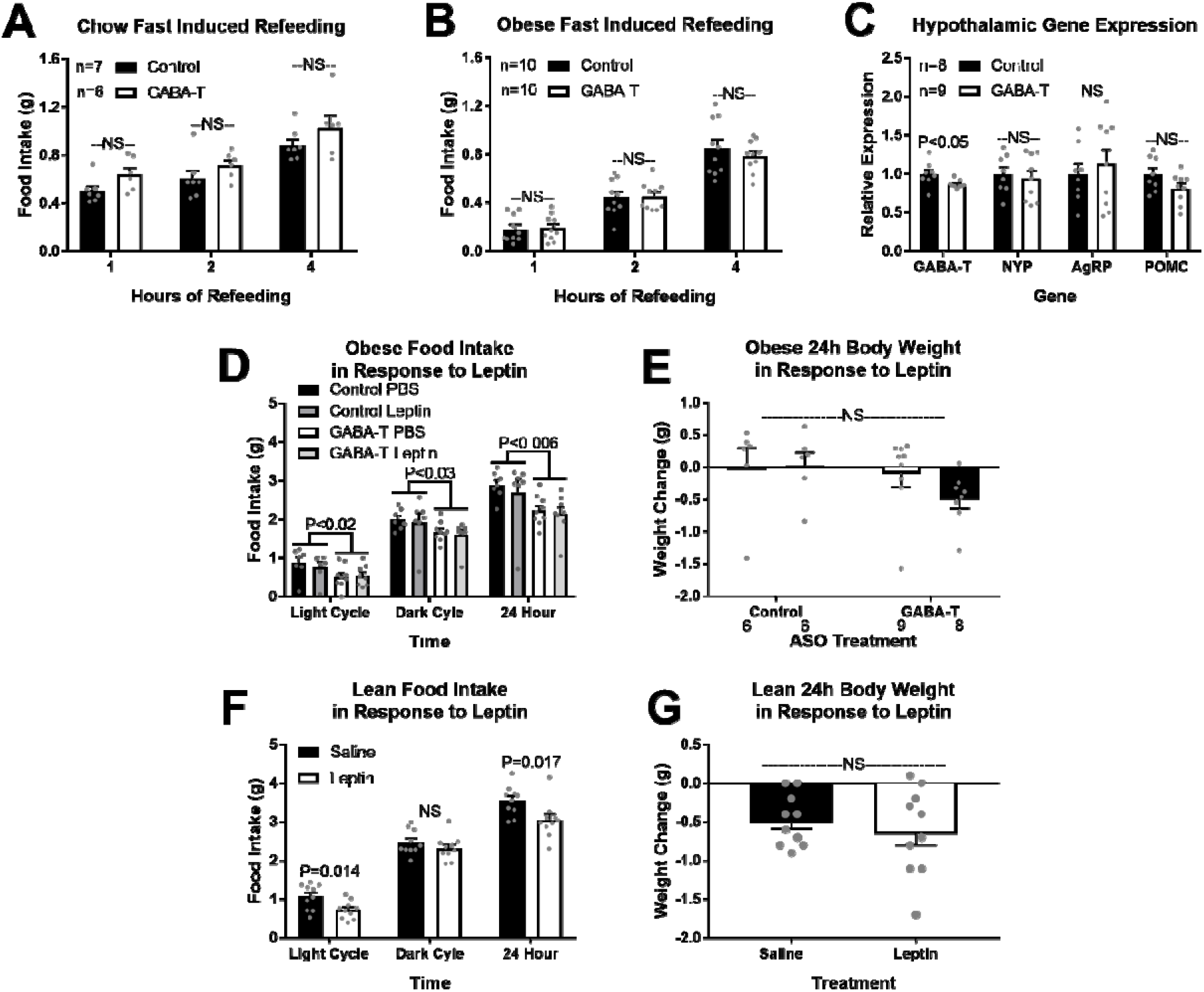
Hepatic GABA-Transaminase knockdown does not affect fast induced refeeding or leptin sensitivity. Refeeding after a 16-hour fast in chow fed lean (A) and diet induced obese (B) mice after 4 weeks of GABA-T targeted or scramble control ASO injections (12.5 mg/kg IP twice weekly). Hypothalamic fasted mRNA expression of GABA-T, neuropeptide Y (NPY), agouti related peptide (AgRP), and pro-opiomelanocortin (POMC; C). The effect of leptin (2 mg/kg IP single injection at 6am) on food intake (D) and body weight change (E) in obese control and GABA-T knockdown mice, and food intake (F) and body weight change (G) in lean mice. All comparisons were made within a timepoint. ^a,b^ bars that do not share a common letter differ significantly (P < 0.05). NS = non-significant. All data are presented as mean ± SEM.

**Extended Data Figure 8.**
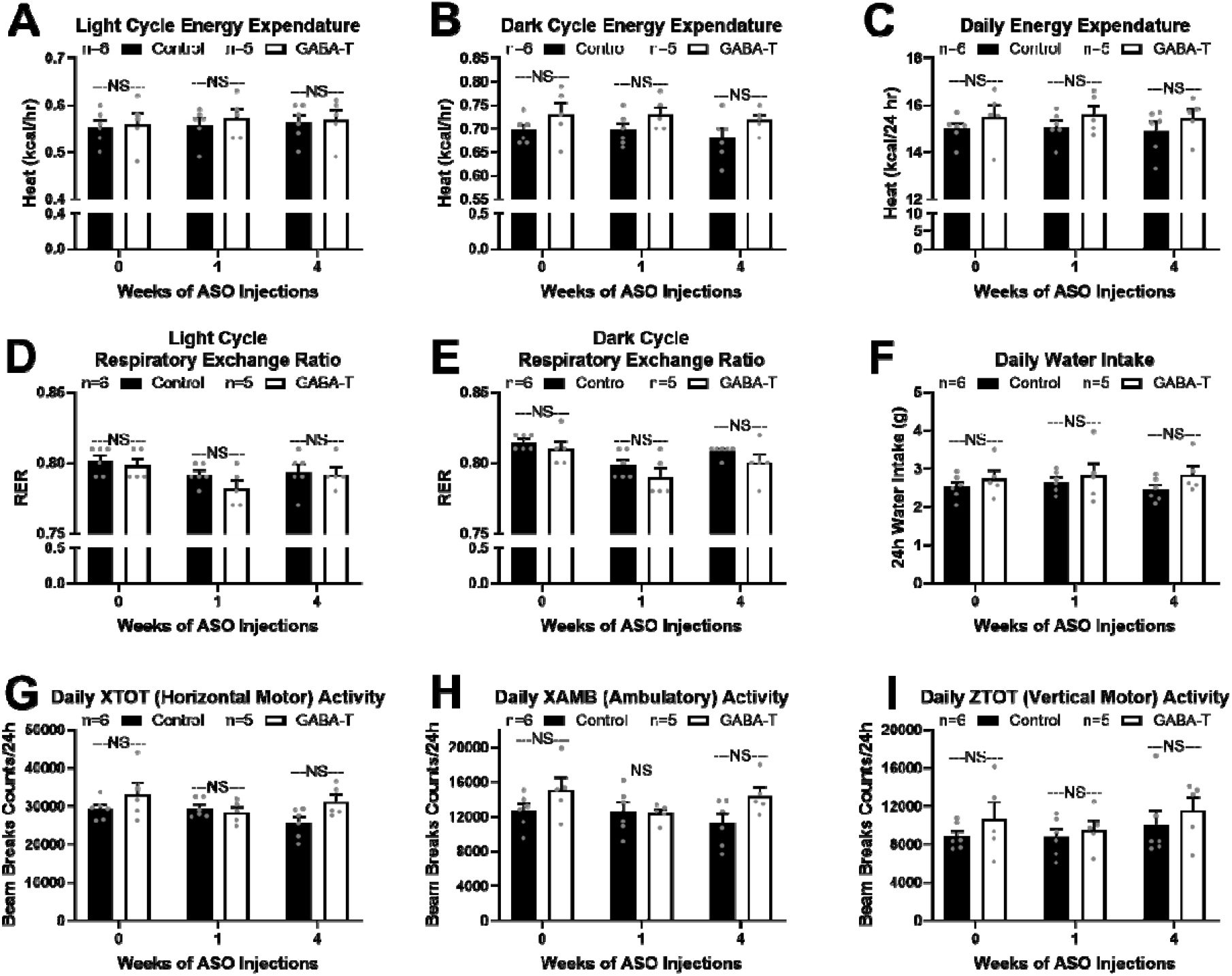
Hepatic GABA-T knockdown does not alter energy expenditure in obesity. Energy expenditure, respiratory exchange ratio, and activity level were assessed by Comprehensive Lab Animal Monitoring System (CLAMS) at the UCDavis Mouse Metabolic Phenotyping Center in diet-induced obese mice after 0, 1, and 4 weeks of GABA-T targeted or scramble control antisense oligonucleotide treatment (ASO; 12.5 mg/kg IP twice weekly). Energy expenditure during the light cycle (A), dark cycle (B), and over 24 hours (C). Respiratory exchange ratio (RER) during the light cycle (D) and dark cycle (E). 24 hour water intake (F). 24 hour activity along the horizontal X axis (XTOT; G), total ambulatory movement (XAMB; H), and vertical Z axis (ZTOT; I). NS = non-significant. All data are presented as mean ± SEM.

**Extended Data Figure 9.**
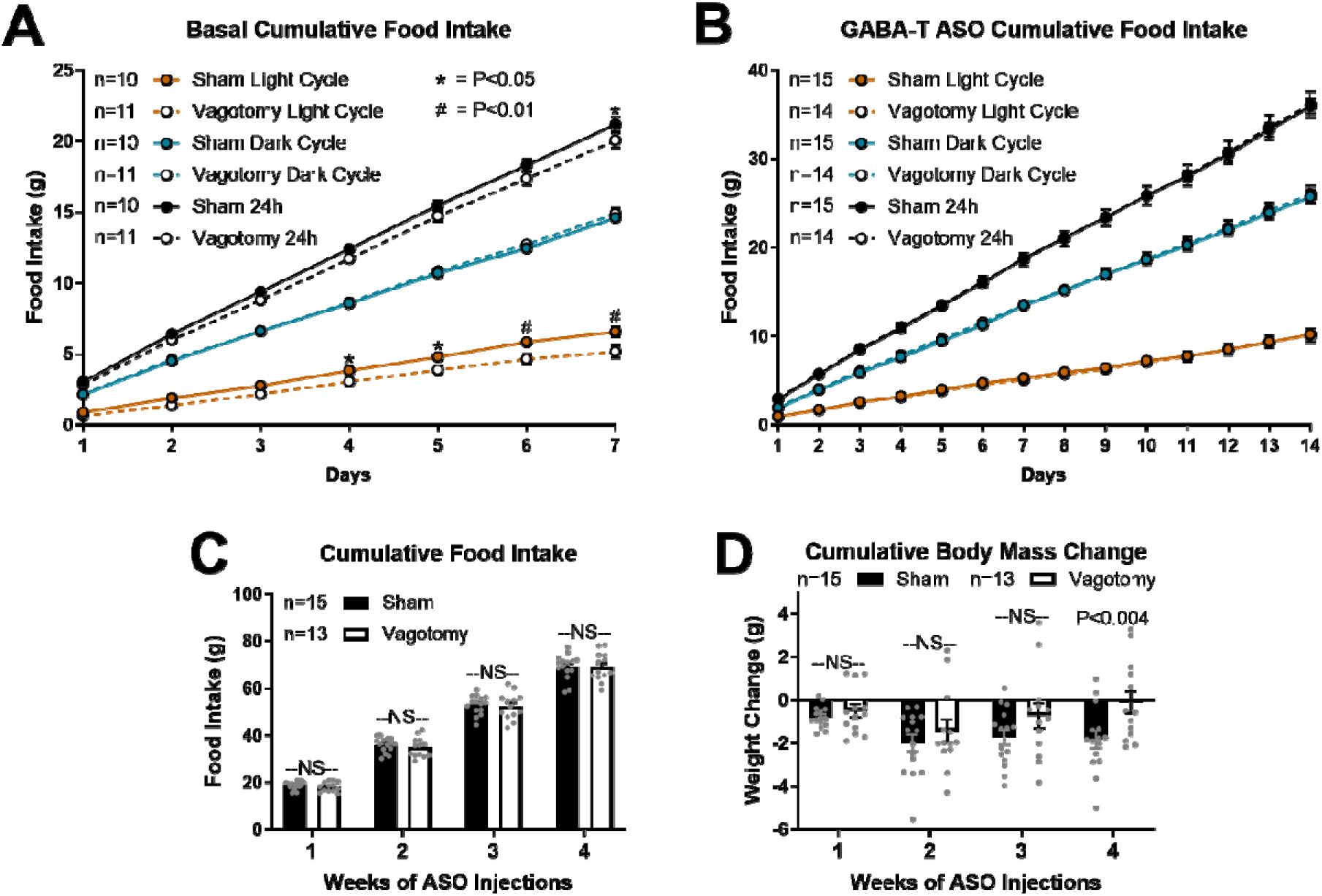
Hepatic vagotomy decreases light cycle food intake on HFD, while GABA-Transaminase knockdown normalizes sham mice food intake to vagotomy mice. Cumulative food intake and body weight in diet-induced obese sham operated and hepatic vagotomized mice during 1 week of baseline feeding (A-B) and during 2 weeks of GABA-T targeted antisense oligonucleotide injections (ASO; 12.5 mg/kg IP twice weekly; C-F). Cumulative basal light cycle, dark cycle, and daily food intake (A) and cumulative body weight change (B). Cumulative ASO light cycle, dark cycle, and daily food intake (C) and cumulative body weight change (D). Weekly cumulative food intake (E) and cumulative body weight change (F). All data are presented as mean ± SEM.

**Extended Data Figure 10.**
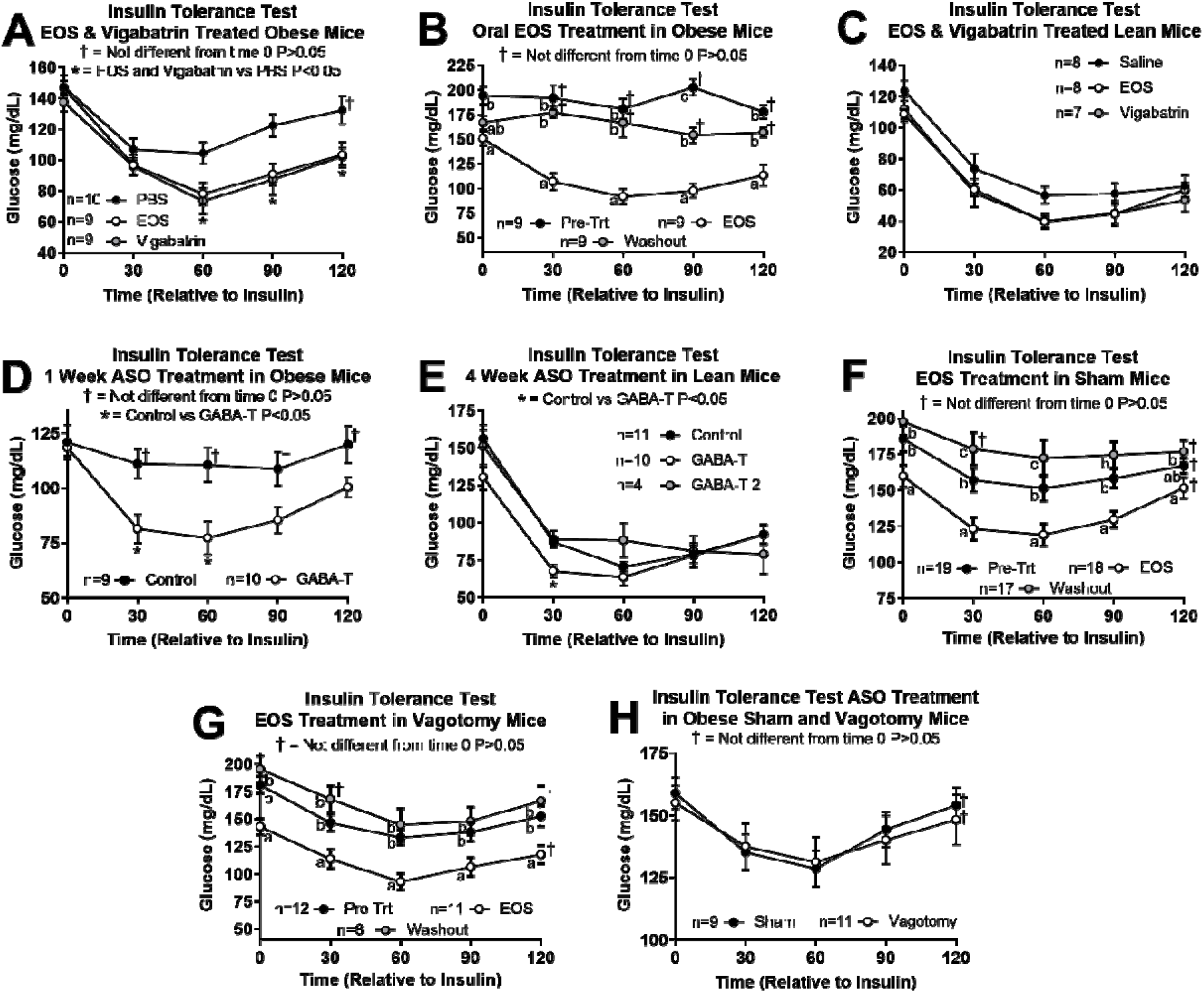
Insulin tolerance tests (ITT) presented as raw glucose values. ITT on day 4 of EOS or Vigabatrin (8 mg/day), or PBS treatment in obese mice (A). ITT pre-treatment, on day 4 of oral EOS (3 g/L in drinking water) treatment, and after a 2-week washout period (B). ITT on day 4 of EOS or Vigabatrin (8mg/day), or PBS treatment in lean mice (C). ITT in obese mice after 1 week of control or GABA-T antisense oligonucleotide (ASO) treatment (D). ITT in lean mice after 4 weeks of control, GABA-T, or GABA-T 2 ASO treatment (E). ITT in sham (F) and vagotomized mice (G) at pre-treatment, on day 5 of EOS (8 mg/day) treatment, and after a 2-week washout period. ITT in obese sham and vagotomized mice after 4 weeks of GABA-T ASO treatment (H). ^a,b,c^ data points that do not share a common letter differ significantly (P < 0.05) within a timepoint. † Denotes the data point is not significantly different from time 0 for that group (P > 0.05). Unless indicated, all other timepoints are significantly different from time 0 within a group of mice. * Denotes significance between groups specified in the panel within a timepoint. All data are presented as mean ± SEM.

## Extended Data Tables and Legends

**Extended Data Table 1.**
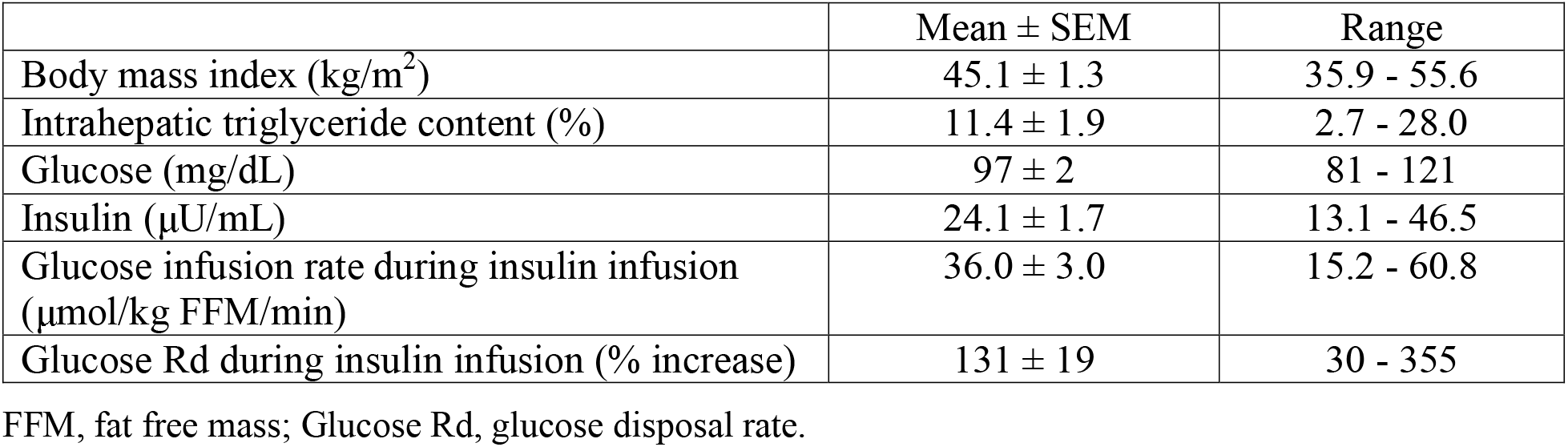
Metabolic characteristics of the study subjects (n=19).

**Extended Data Table 2.** Regression coefficient estimates showing the association between hepatic mRNA expression of genes involved in GABA production (*ABAT*) and GABA transport (*SLC6A6, SLC6A8, SLC6A12, and SLC6A13*) and basal plasma insulin concentration (μU/mL) or hepatic insulin sensitivity index (HISI).

| Basal Plasma Insulin Concentration (μU/mL) | | | | | |
| --- | --- | --- | --- | --- | --- |
|  | Estimate | SEM | Lower CI | Upper CI | P- Value |
| **Intercept** | -52.50 | 40.31 | -143.67 | 38.68 | 0.2251 |
| **IHTG (%)** | 0.30 | 0.14 | -0.01 | 0.62 | 0.0577 |
| ***ABAT*** | 18.14 | 5.28 | 6.19 | 30.09 | 0.0075 |
| ***SLC6A12*** | -14.71 | 4.29 | -24.41 | -5.01 | 0.0075 |
| ***SLC6A13*** | 1.90 | 3.14 | -5.19 | 8.99 | 0.5595 |
| ***SLC6A6*** | 3.02 | 2.40 | -2.41 | 8.45 | 0.2395 |
| ***SLC6A8*** | -1.87 | 1.47 | -5.18 | 1.45 | 0.2342 |
| HISI |  |  |  |  |  |
|  | Estimate | SEM | Lower CI | Upper CI | P- Value |
| **Intercept** | 16.41 | 7.23 | 0.06 | 32.76 | 0.0493 |
| **IHTG (%)** | -0.05 | 0.03 | -0.11 | 0.00 | 0.0674 |
| ***ABAT*** | -2.94 | 0.95 | -5.08 | -0.80 | 0.0127 |
| ***SLC6A12*** | 2.38 | 0.77 | 0.64 | 4.12 | 0.0127 |
| ***SLC6A13*** | -0.71 | 0.56 | -1.98 | 0.56 | 0.2379 |
| ***SLC6A6*** | -0.21 | 0.43 | -1.19 | 0.76 | 0.6340 |
| ***SLC6A8*** | 0.17 | 0.26 | -0.43 | 0.76 | 0.5416 |

**Extended Data Table 3.**
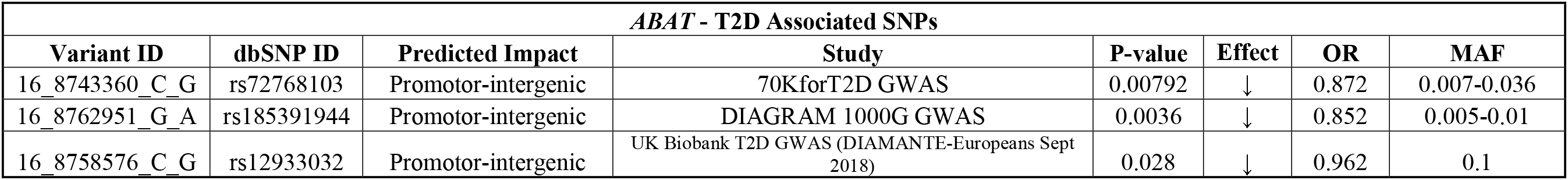
Single Nucleotide Polymorphisms (SNPs) in the promoter of the ABAT gene, which encodes for GABA transaminase, are associated with a decreased odds ratio (OR) for type 2 diabetes (T2D; Source: knowledge portal diabetes database). MAF – minor allele frequency.

**Extended Data Table 4.** Single Nucleotide Polymorphisms (SNPs) that result in missense mutations in GABA transporters are associated with BMI (Source: knowledge portal diabetes database). MAF – minor allele frequency.

| *SLC6A12* - BMI Associated SNPs |  |  |  |  |  |  |  |
| --- | --- | --- | --- | --- | --- | --- | --- |
| Variant ID | dbSNP ID | Predicted Impact | Study | P-value | Effect | Effect Size | MAF |
| 12_313824_G_A | rs199521597 | Missense: Replaces Alanine with Valine | GIANT 2018 BMI, Height exome chip analysis: African Americans | 0.0026 | ↑ | 3 | Not Reported |
| 12_319125_A_G | rs557881 | Missense: Replaces Cysteine with Arginine | GIANT UK Biobank GWAS | 0.0031 | ↑ | 0.0053 | 0.426 |
| 12_309921_T_C | rs143648821 | Missense: Replaces Isoleucine with Valine | FUSION exome chip analysis | 0.00316 | ↑ | 0.815 | 0.00105 |
| 12_309864_C_T | rs11061915 | Missense: Replaces Valine with Isoleucine | 13K exome sequence analysis | 0.00623 | ↑ | 0.299 | 0.00354 |
| 12_300248_C_G,T | rs147574089 | Missense: Replaces Glutamate with Glutamine | GIANT 2018 BMI, Height exome chip analysis: Hispanics | 0.0081 | ↑ | 0.65 | NaN |
| 12_300298_C_T | rs537332809 | Missense: Replaces Arginine with Glutamine | 13K exome sequence analysis | 0.0141 | ↑ | 1.09 | 0.000381 |
| *SLC6A6* - BMI Associated SNPs |  |  |  |  |  |  |  |
| Variant ID | dbSNP ID | Predicted Impact | Study | P-value | Effect | Effect Size | MAF |
| 3_14523209_G_A | rs141254266 | Missense: Replaces Valine with Isoleucine | 13K exome sequence analysis | 0.00922 | ↑ | 0.866 | 0.00009-0.0003 |
| 3_14523296_G_A | rs41284017 | Missense: Replaces Valine with Isoleucine | GIANT 2018 BMI, Height exome chip analysis | 0.016 | ↑ | 0.028 | 0.004-0.0165 |
| 3_14526454_G_A | rs200063855 | Missense: Replaces Arginine with Histadine | GIANT 2018 BMI, Height exome chip analysis: South Asians | 0.033 | ↓ | -0.71 | 0.00007-0.0002 |
| *SLC6A13* - BMI Associated SNPs |  |  |  |  |  |  |  |
| Variant ID | dbSNP ID | Predicted Impact | Study | P-value | Effect | Effect Size | MAF |
| 12_332337_C_T | rs202217743 | Missense: Replaces Valine with Leucine, or Methionine | FinnMetSeq exome sequence analysis | 0.0123 | ↑ | 1.26 | 0.0001 |
| 12_331781_C_T,G | rs147388541 | Missense: Replaces Aspartate with Histadine, or Asparagine | GIANT 2018 BMI, Height exome chip analysis: South Asians | 0.015 | ↑ | 0.59 | 0.003-0.02 |
| 12_347102_C_T | rs138506621 | Missense: Replaces Glutamate with Lysine | GIANT 2018 BMI, Height exome chip analysis: East Asians | 0.034 | ↑ | 2.1 | 0.00014-0.001 |
| 12_352884_C_T | rs543043546 | Missense: Replaces Glycine with Serine | 13K exome sequence analysis | 0.0341 | ↑ | 0.489 | 0.00001-0.00055 |

## Materials and Methods

### Animals

All studies were conducted using male wildtype C57BL/6J mice purchased from Jackson Laboratories or bred in-house (Bar Harbor, ME). Mice were kept on a 14-hour light/10-hour dark schedule and housed 3-5 mice per cage until 1 week prior to study initiation, at which point animals were individually housed. We conducted studies in lean chow fed mice (7013 NIH-31, Teklad WI, 3.1 kcal/g, 18% kcal from fat, 59% kcal from carbohydrate, 23% kcal from protein) at 12-16 weeks of age. Studies in diet-induced obese mice dosed intraperitoneally with GABA transaminase inhibitors, mice treated with ethanolamine-O-sulfate in their drinking water, and mice treated with GABA-T targeted or scramble control antisense oligonucleotides (ASO) were performed after 8-10 weeks on a high fat diet (TD 06414, Teklad WI, 5.1 kcal/g, 60.3% kcal from fat, 21.3% kcal from carbohydrate, 18.4% kcal from protein; 20-26 weeks of age). For ASO studies, mice were stratified by body weight and assigned to an injection treatment (control or GABA-T). Studies in obese vagotomy and sham mice were performed after 9 weeks of high fat diet feeding. Unless fasted, mice had *ad libitum* access to food and water. All studies were approved by The University of Arizona Institutional Animal Care and Use Committee.

### Antisense Oligonucleotide Studies

Wildtype lean or diet-induced obese mice received twice-weekly intraperitoneal injections (12.5 mg/kg; 0.1 mL/10 g body weight) of murine GABA-Transaminase (GABA-T) targeted antisense oligonucleotides (ASO; IONIS 1160575; 5′-AAGCTATGGACTCGGT-3′) or scramble control ASO (IONIS 549144; 5′-GGCCAATACGCCGTCA-3′) for 1 or 4 weeks prior to experimentation. The control ASO does not have complementarity to known genes and was employed to demonstrate the specificity of target reduction. 2’, 4’-constrained 2’-O-ethyl (cEt) ASOs were synthesized at Ionis Pharmaceuticals (Carlsbad, CA) as described previously ^45^. Treatment with this GABA-T targeted ASO reduces hepatic GABA-T mRNA by 98% in adult mice within 1 week (Fig. 2A). For studies after 1 week of ASO injections, we performed an OGTT (day 8), ITT (day 10), and mice were sacrificed to assess liver slice GABA release (day 12). For studies after 4 weeks of ASO injections, we performed an OGTT (day 29), ITT (day 30), and completed all additional studies within 1 week while continuing biweekly ASO injections before mice were sacrificed to assess liver slice GABA release.

### Hepatic Vagotomy Surgeries

Surgeries were performed in 12-week old male C57BL/6J mice under isoflurane anesthesia. Mice were randomly assigned to a surgical group (sham or vagotomy). A ventral midline incision through the skin and peritoneum allowed us to isolate the hepatic vagus nerve as it branched from the esophagus. In vagotomized mice, we severed the hepatic vagal nerve, while it remained intact in sham operated mice. The peritoneum was sutured with absorbable polyglactin 910 suture and the skin with nylon suture. Mice were given a single post-operative dose of slow release formulated buprenorphine analgesic (1.2 mg/kg slow release, sub-cutaneous). We monitored food intake and body weight daily and removed sutures 7 days post-operation.

### GABA Transaminase Inhibitor Studies

Wildtype lean and obese mice were randomly divided into treatment groups and dosed daily with 8 mg of ethanolamine-O-sulfate (EOS; Sigma-Aldrich, St. Louis, MO), vigabatrin (United States Pharmacopeia, Rockville, MD) or PBS. Obese sham and vagotomy mice were dosed daily with 8 mg of EOS. In all cases, basal bleeds were taken prior to initiation of an ITT. Lean mice received treatment by oral gavage (0.3 mL/mouse) while obese mice were treated by intraperitoneal injection (0.3 mL/mouse). Pre-treatment studies were conducted in the week immediately prior to beginning drug administration. Daily doses took place at 9 am each day and oral glucose tolerance tests (OGTT) and insulin tolerance tests (ITT) were performed on the third and fourth days of treatment in lean mice, respectively. In wildtype obese mice, OGTT and ITT were performed on the fourth day of treatment in separate cohorts. On the fifth day of treatment we preformed 2DG clearance studies. In sham/vagotomy mice, OGTT and ITT were performed on the fourth and fifth day of treatment, respectively. After a 2-week washout period without treatment injection, a basal bleed and bleed 15 minutes after an oral glucose gavage (2.5 mg/kg) to determine oral glucose stimulated insulin secretion were performed.

In a separate set of studies, an ITT was performed in obese mice to establish insulin resistance. Subsequently, EOS was provided *ad libitum* in the water (3 g/L) for 4 days. An OGTT and ITT were performed on days 3 and 4 of treatment, respectively. The water was then removed, and an ITT was performed 2 weeks later to establish the timing of restoration of insulin resistance after drug removal.

### Food Intake Studies

We measured food weight and body weight at 6am and 6pm for the first 14 days of control or GABA-T ASO injections in lean and obese wildtype mice (Fig. 6) and GABA-T ASO injections in sham and vagotomy mice (Fig. 7). Weeks 3 and 4 food and body weight measurements were taken at 6pm on day 21 and 28 of ASO injections.

### Fast-Induced Refeeding

Mice were housed on wood chip bedding (Harlan Laboratories; Cat #7090 Sani-Chips) to limit consumption of nutrients from bedding during the fasting period. Mice were fasted for 16 hours beginning at 5pm and food was returned at 9am. Food and body weight were measured at 10am, 11am, and 1pm to determine 1, 2, and 3-4 hour fast-induced refeeding.

### Leptin Food Intake Study

Lean mice or diet-induced obese control or GABA-T targeted ASO treated mice received an intraperitoneal injection of phosphate buffered saline (PBS; 0.1 mL/10 g body weight) on day 1 and leptin (2 mg/kg; 0.1 mL/10 g body weight; CAT# 498-OB, R&D Systems, Minneapolis, MN) on day 2 at 6am. Mice were not fasted before injections. Food and body weight were measured every day at 6am and 6pm.

### ASO Energy Expenditure and Body Composition

Energy expenditure studies and body composition measurements were performed by the UC Davis Mouse Metabolic Phenotyping Center. Twelve male DIO (C57BL/6J, strain 380050) mice were sent to UC Davis from the Renquist Lab at ~20 weeks of age and were acclimated for 2 weeks on investigator provided diet (Research diets D12492). Mice were injected IP with control or ABAT ASO (12.5 mg/kg) twice a week for 4 weeks (8 injections total). Energy expenditure and physical activity were evaluated in 6 saline and 6 ABAT ASO treated mice by indirect calorimetry in the CLAMS (Comprehensive Lab Animal Monitoring System, Columbus Instruments) system three times at baseline (−3), 7, and 28 days post first ASO injection. Body composition was assessed under isoflurane anesthesia by DEXA after each CLAMS run. Animals were allowed to recover from anesthesia before returning to their home cage except for the final (3^rd^) DEXA run, after which animals were terminated and no tissues collected. Animals were acclimated to the CLAMS cages for 48 hours and to the light and temperature-controlled chamber for 24 hours prior to testing. Animals were held and calorimetry data was collected for 48 hrs. Analyzed data constitutes data collected from 48 hours of continuous measurement (2 light/2 dark cycles). Oxygen consumption and carbon dioxide production were measured and used to calculate energy expenditure (or heat production, kilocalories (kcal)) and respiratory exchange ratio (RER: VCO2/VO2). Cage-mounted sensors detected and recorded measurements of physical activity, food intake and water intake. Body composition was measured by dual-energy X-ray absorptiometry under isoflurane anesthesia, using a Lunar PIXImus II Densitometer (GE Medical Systems, Chalfont St. Giles, UK) immediately after completion of the indirect respiration calorimetry measurements.

### EOS Body Composition

Body composition was assessed in diet-induced obese mice on day 0 and 7 days after continuous provision of EOS in the drinking water (3 g/L) using an EchoMRI 900 with A10 insert for mice. We performed calibration daily and had the water stage set to on.

### Hyperinsulinemic Euglycemic Clamps

Clamps were performed as previously described ^46^. Briefly, one week prior to the experiment, diet-induced obese mice underwent surgical catheterization of the jugular vein under isoflurane anesthesia. Mice were given a single post-operative dose of slow release formulated buprenorphine analgesic (1.2 mg/kg slow release, sub-cutaneous) and food and body weight were assessed daily for 1 week. The same day following surgery completion mice received their first injection of either the scramble control or GABA-T targeted ASO (12.5 mg/kg; 0.1 mL/10 g body weight). Four days post-surgery mice received a second ASO injection, and clamps were performed 7 days after catheterization. Following a 5-h fast (starting at 0800h), clamps were performed using unrestrained, conscious mice. The clamp procedure consisted of a 90-min tracer equilibration period (−90 to 0 min), followed by a 120-min clamp period (0 to 120 min). To begin the equilibration period, mice were infused with 3-[^3^H]-D-glucose (0.05 μCi/μl in saline) at a rate of 10 μL/min for 2 minutes and then decreased to a rate of 1 μL/min for the remaining 90 minutes. All blood was collected from the tip of the tail in heparinized capillary tubes and immediately spun down to collect plasma for tracer and insulin analysis. All timepoints during the clamp are relative to the start of insulin infusion at time 0. Blood for tracer and insulin analysis was taken at −10 and 0 minutes. At 0 minutes mice were infused with donor blood (5 μL/min continuously throughout study), insulin (4 mU/kg/min; pump rate 1 μL/min continuously throughout study), and a variable infusion of 3-[^3^H]-D-glucose (0.05 μCi/μl in 50% dextrose) to maintain euglycemia. Glucose was assessed from whole-blood directly from the tail tip by glucometer (Manufacture # D2ASCCONKIT, Bayer, Leverkusen, Germany) every 10 minutes for 120 minutes starting at time 0. The glucose infusion rate was adjusted to maintain a blood glucose concentration in the range of 100-120 mg/dL. Blood for tracer measurements was taken at 80, 90, 100, and 120 minutes, and blood for insulin was taken at 100 and 120 minutes. To estimate insulin-stimulated glucose fluxes in tissues, mice were given a bolus of ^14^C-2-deoxyglucose (12 μCi in 48 μL followed by a 100 μL saline flush) after the 120min timepoint. Blood samples were collected 2, 10, and 25 minutes following the bolus. Mice were anesthetized with isoflurane using the bell-jar method and the soleus, quadricep, calf, white adipose tissue, and perirenal adipose tissue were collected and immediately frozen in liquid nitrogen for analysis of tissue specific glucose uptake. Plasma samples were deproteinized using barium hydroxide and zinc sulfate and tracer analysis and tracer standards was processed as described ^46^. Plasma glucose was analyzed by colorimetric assay (Cat. # G7519, Pointe Scientific Inc., Canton MI) from the samples collected for tracer measurement. Plasma insulin was analyzed by enzyme-linked immunosorbent assay (ELISA; Cat. # 80-INSMSU-E01,E10, Alpco, Salem, NH).

### Oral Glucose Tolerance Test

Oral glucose (2.5 g/kg; 0.1 mL/10 g body weight; Chem-Impex Int’l Inc., Wood Dale, IL) was given to 4 hour fasted individually housed mice. All glucose tolerance tests began at 1 pm and glucose was measured in whole blood, collected from the tail vein, by glucometer (Manufacture # D2ASCCONKIT, Bayer, Leverkusen, Germany) at 0, 15, 30, 60, 90, and 120 minutes after glucose gavage. Blood for serum insulin (oral glucose stimulated insulin secretion; OGSIS) and glucose determination was collected from the tail vein 15 minutes following glucose administration.

### Insulin Tolerance Test

Intraperitoneal insulin (0.5 U/kg; 0.1 mL/10 g body weight; Sigma Aldrich, St. Louis, MO) was given to 4 hour fasted individually housed mice. All insulin tolerance tests began at 1 pm and glucose was measured in whole blood, collected from the tail vein, by glucometer (Manufacture # D2ASCCONKIT, Bayer, Leverkusen, Germany) at 0, 30, 60, 90, and 120 minutes after insulin injection.

### Liver Slice Explant Studies

Liver slices from ASO treated mice were incubated *ex vivo* to measure hepatic GABA release. A peristaltic pump perfusion system was used to deliver warmed KH buffer to the liver through the portal vein. Briefly, mice were anesthetized with an intraperitoneal injection of ketamine (10 mg/mL) and diazepam (0.5 mg/mL). Once mice were unresponsive, an incision in the lower abdomen through the skin and peritoneal membrane was made vertically through the chest along with transverse incisions on both sides to expose the liver. A 30-gauge needle was inserted into the hepatoportal vein to blanch the liver. The inferior vena cava was cut to relieve pressure in the circulatory system and allow blood to drain. The perfusion continued for several minutes at a rate of 4 mL/minute until the liver was completely blanched. The liver was removed and washed in warm PBS before being sliced into 0.2 mm slices using a Thomas Sadie-Riggs Tissue Slicer. Two liver slices were taken from each mouse. Tissue slices were placed individually into a well on a 12-well plate pre-filled with 1mL of KH buffer that had been sitting in an incubator set to 37°C and gassed with 5% CO_2_. Liver slices were incubated in the initial well for 1 hour to stabilize before being transferred to a fresh well pre-filled with KH buffer. After 1 hour in the second well, tissue and media were collected. Liver slice samples and KH media samples from both wells of each mouse were pooled. Liver slices were snap frozen in liquid nitrogen, while media was centrifuged for 5 minutes at 10,000xg at 4°C to remove tissue debris and both were frozen and stored at −80°C pending analysis.

### GABA Analysis

For all liver slice GABA release data, we thawed the media collected from the *ex vivo* hepatic slice culture on ice. We then measured GABA in the supernatant using a commercially available ELISA (REF# BA E-2500, Labor Diagnostika Nord, Nordhorn, Germany). μmol GABA concentrations were corrected for liver slice DNA concentrations.

### ^3^H-2-deoxy-D-glucose Uptake Studies

On the fifth day of EOS or PBS treatment ^3^H-2-deoxy-D-glucose (2DG; 10 uCi/mouse; PerkinElmer, Waltham, MA) was given to 4-hour fasted individually housed mice. Studies were performed in 2 cohorts on 2 different days. 2DG was given by oral gavage in a solution of glucose (2.5 g/kg) and each mouse received the same dose based off the average body weight for their cohort (0.1 mL/10 g body weight). Mice were anesthetized by isoflurane and sacrificed by cervical dislocation 45 minutes following oral gavage. Liver, soleus, quadriceps femoris, and gonadal white adipose tissue were collected, weighed, and dissolved overnight in 1N NaOH (0.5 mL/50 mg tissue) at 55°C on a shaker plate. 0.5 mL of dissolved tissue was added to 5 mL of scintillation cocktail (Ultima Gold, PerkinElmer, Waltham, MA) and disintegrations per minute (DPM) were measured using a LS 6500 Multipurpose Scintillation Counter (Beckman Coulter, Brea, CA). DPM/g tissue weight was determined for each tissue and normalized based on a correction factor calculated by the average total DPM/g for all tissues divided by the total DPM/g for all tissues of the individual mouse.

### cGMP Analysis

On the fifth day of EOS or PBS treatment, mice were sacrificed and the quadricep and soleus tissues were collected and frozen on dry ice. Prior to analysis, frozen quadricep were powdered using a liquid nitrogen cooled mortar and pestle to obtain homogenous muscle samples. 15-20 mg of quadricep and the entire soleus tissue (6-12 mg) were homogenized in 200 μL of a 5% trichloroacetic acid solution. Following 15 minutes of centrifugation at 3,000xg at 4°C, supernatant was transferred to a fresh tube for analysis of muscle cGMP by enzyme-linked immunosorbent assay (ELISA; ADI-900-164, Enzo Life Sciences, Farmingdale, NY).

### Liver Analysis

Prior to analysis, frozen livers were powdered using a liquid nitrogen cooled mortar and pestle to obtain homogenous liver samples. To measure liver DNA content (ng dsDNA/g tissue), 10-20 mg of powdered liver was sonicated in 500 μL DEPC H_2_O and dsDNA determined by fluorometric assay (Cat. # P7589, Invitrogen, Waltham, MA). Whole liver and hypothalamic mRNA were isolated from powered liver samples with TRI Reagent® (Life Technologies, Grand Island, NY) and phenol contamination was eliminated by using water-saturated butanol and ether as previously described ^47^. cDNA was synthesized by reverse transcription with Verso cDNA synthesis kit (Thermo Scientific, Inc., Waltham, MA), and qPCR performed using SYBR 2X mastermix (Bio-Rad Laboratories, Hercules, CA) and the Biorad iQ™5 iCycler (Bio-Rad Laboratories, Hercules, CA). Expression of GABA-Transaminase (ABAT), β-actin (ACTβ), insulin (Ins), neuropeptide Y (NPY), agouti related peptide (AgRP), and pro-opiomelanocortin (POMC) mRNA were measured using the following primers (5’→3’): ABAT, Forward – CTGAACACAATCCAGAATGCAGA Reverse – GGTTGTAACCTATGGGCACAG; ACTβ, Forward – TCGGTGACATCAAAGAGAAG Reverse – GATGCCACAGGATTCCATA; Ins, Forward – GCTTCTTCTACACACCCATGTC Reverse – AGCACTGATCTACAATGCCAC; NPY, Forward – CTCGTGTGTTTGGGCATTCT Reverse – CTTGCCATATCTCTGTCTGGTG; AgRP, Forward – GTACGGCCACGAACCTCTGT Reverse – TCCCCTGCCTTTGCCCAA; POMC, Forward – GGTGAAGGTGTACCCCAACGT Reverse – GACCTGGCTCCAAGCCTAATGG. LinReg PCR analysis software was used to determine the efficiency of amplification from raw output data ^48^. ACTβ served as the reference gene for liver and brain tissue, and Ins served as the reference gene for pancreas tissue for calculating fold change in ABAT gene expression using the efficiency^−ΔΔCt^ method ^49^.

### Serum Assays

Within 2 hours of collection, blood was left to clot at room temperature for 20 minutes. Thereafter, the blood was centrifuged at 3,000xg for 30 minutes at 4°C and serum was collected. Serum was stored at −80°C until metabolite and hormone analyses. We used a colorimetric assay to analyze serum glucose (Cat. # G7519, Pointe Scientific Inc., Canton MI). Serum insulin was analyzed by enzyme-linked immunosorbent assay (ELISA; Cat. # 80-INSMSU-E01,E10, Alpco, Salem, NH). Serum glucagon was analyzed by enzyme-linked immunosorbent assay (ELISA; Cat. # 10-1281-01, Mercodia, Uppsala, Sweden) from tail vein blood collected at 9 am from fed mice.

### Studies Conducted in Human Subjects

A total of 19 men and women with obesity who were scheduled for bariatric surgery at Barnes-Jewish Hospital in St. Louis, MO participated in this study, which was conducted at Washington University School of Medicine in St. Louis. Subjects provided written, informed consent before participating in this study, which was approved by the Human Research Protection Office at Washington University School of Medicine in St. Louis, MO (ClinicalTrials.gov NCT00981500). Intrahepatic triglyceride content was determined by using magnetic resonance imaging (3.0-T superconducting magnet; Siemens, Iselin, NJ) in the Center for Clinical Imaging Research. A 7-hour (3.5-h basal period and 3.5-h insulin infusion period) HECP, in conjunction with stable isotopically labelled glucose tracer infusion, was then conducted in the Clinical Translational Research Unit (CTRU), as previously described ^50^. This procedure was performed to determine: i) hepatic insulin sensitivity, which was assessed as the product of the basal endogenous glucose production rate (in μmol·kg fat-free mass (FFM)^−1^·min^−1^) and fasting plasma insulin concentration (in mU/L).

#### Liver RNA sequencing (RNA-seq)

Liver tissue was obtained by needle biopsy during the bariatric surgical procedure, before any intra-operative procedures were performed. Liver tissue was rinsed in sterile saline, immediately frozen in liquid nitrogen, then stored at −80°C until RNA extraction. Total RNA was isolated from frozen hepatic tissue samples by using Trizol reagent (Invitrogen, Carlsbad, CA). Library preparation was performed with total RNA and cDNA fragments were sequenced on an Illumina HiSeq-4000. The fragments per kilobase million reads upper quartile (FPKM-UQ) values were calculated and used for further gene expression analyses. All RNA-seq data used in this study have been deposited into the NCBI GEO database under accession number GSE144414.

### Statistics

We analyzed the data in SAS Enterprise Guide 7.1 (SAS Inst., Cary, NC), using a mixed-model ANOVA for all analyses except the multivariate regression analyses performed on human clinical data. ANOVA tests do not have a one-tailed vs. two-tailed option, because the distributions they are based on have only one tail. When comparisons between all means were required, we used a Tukey’s adjustment for multiple comparisons. When comparisons of means were limited (e.g. only within a timepoint or treatment), we used a Bonferonni correction for multiple comparisons. For the analysis of ITT and OGTT, repeated measures ANOVA were performed by including time point in the analysis. Analyses were conducted separately for chow and HFD fed mice. For the studies using the GABA-T inhibitors the main effect was treatment (PBS, Vigabatrin, or EOS). For EOS treated sham and vagotomy mice the main effects were surgery (sham or vagotomy) and treatment (PBS or EOS). Pre-, during, and post-treatment measures were taken for basal glucose, insulin, and glucose stimulated insulin, thus a repeated measure analysis including time (pre-, during, or post-treatment) was performed separately within each injection or surgical treatment. For analysis of the effect of EOS on 2DG uptake and cGMP, analysis was performed separately for each tissue. For ASO studies the main effect was injection treatment (control of GABA-T ASO). For the vagotomy analyses the main effect was surgery (sham or vagotomy). Pre-treatment and ASO week 4 measures were taken for basal glucose, insulin, and the glucose:insulin ratio, thus a repeated measure analysis including time (pre-, or week 4-treatment) was performed separately within each injection or surgical treatment. A multivariate regression analysis was performed on data from human clinical patients using IHTG%, ABAT mRNA, and SLC6a12 mRNA as explanatory variables for variations in serum insulin, HOMA-IR, M-Value, and Glucose Rd. Statistics performed by the UC Davis Mouse Metabolic Phenotyping Center are described as follows: data are presented as means ± SEM and a Student t-test was used to test for significant differences between groups. Multiple linear regression analysis (analysis of covariance, ANCOVA) was used to assess the impact of covariates (e.g. body weight or lean mass) on energy expenditure. The EE ANCOVA analysis done for this work was provided by the NIDDK Mouse Metabolic Phenotyping Centers (MMPC, www.mmpc.org) using their Energy Expenditure Analysis page (http://www.mmpc.org/shared/regression.aspx) and supported by grant DK076169. All insulin tolerance tests are presented as a percentage of baseline glucose in main and Extended Data figures, and additionally presented as raw glucose values in Extended Data Fig. 10. Human data was analyzed using a multivariate regression including IHTG content, and *ABAT, SLC6A6, SLC6A8, SLC6A12,* and *SLC6A13* mRNA expression as independent variables with Type 3 test of fixed effects used to determine significance and estimates derived from maximum likelihood estimation. All graphs were generated using GraphPad Prism 8 (GraphPad Software Inc., La Jolla, CA).

## Data Availability

The datasets generated and/or analyzed during the current study are available in the Mendeley data repository at a link provided in the cover letter. Datasets will be made public upon acceptance of this manuscript.

## Endnotes

## Acknowledgments

The authors wish to thank Drs. Richard Lee and Mark Graham of Ionis Pharmaceuticals, Carlsbad, CA for providing both the GABA-T and control antisense oligonucleotides, and the UC Davis MMPC Energy Balance, Exercise, & Behavior Core (NIH grant U24-DK092993) for performing energy expenditure and physical activity measurements by indirect respiratory calorimetry using CLAMS and assessing body composition by DEXA.

## Author Contributions

CEG – Experimental design and project conceptualization, performed experiments and wet lab analyses, wrote initial draft of manuscript, generated figures, reviewed and edited manuscript.

SG – Performed experiments and wet lab analyses, reviewed and edited manuscript.

SMB – Performed experiments and wet lab analyses, reviewed and edited manuscript.

KM – Performed SNP analysis, reviewed and edited manuscript.

JMK – Performed experiments and wet lab analyses, reviewed and edited manuscript.

SNW – Performed clamp surgeries and experiments, reviewed and edited manuscript.

FAD – Performed clamp surgeries and experiments, reviewed and edited manuscript.

JY – Conducted human clinical trial, reviewed and edited manuscript.

SK – Conducted human clinical trial, reviewed and edited manuscript.

BJR – Experimental design and project conceptualization, performed surgeries, analyzed statistics, reviewed and edited manuscript.

## Author Information

Reprints and permissions information is available at www.nature.com/reprints. Competing Interests: The results presented in this paper have resulted in patent cooperation treaty Application No. 62/511,753 and 62/647,468: METHODS AND COMPOSITIONS FOR REGULATING GLUCOSE HOMEOSTASIS. Correspondence and requests for materials should be addressed to

